# A high resolution single molecule sequencing-based Arabidopsis transcriptome using novel methods of Iso-seq analysis

**DOI:** 10.1101/2021.09.02.458763

**Authors:** Runxuan Zhang, Richard Kuo, Max Coulter, Cristiane P. G. Calixto, Juan Carlos Entizne, Wenbin Guo, Yamile Marquez, Linda Milne, Stefan Riegler, Akihiro Matsui, Maho Tanaka, Sarah Harvey, Yubang Gao, Theresa Wießner-Kroh, Martin Crespi, Katherine Denby, Asa ben Hur, Enamul Huq, Michael Jantsch, Artur Jarmolowski, Tino Koester, Sascha Laubinger, Qingshun Quinn Li, Lianfeng Gu, Motoaki Seki, Dorothee Staiger, Ramanjulu Sunkar, Zofia Szweykowska-Kulinska, Shih-Long Tu, Andreas Wachter, Robbie Waugh, Liming Xiong, Xiao-Ning Zhang, Anireddy S.N. Reddy, Andrea Barta, Maria Kalyna, John WS Brown

## Abstract

**Background:** Accurate and comprehensive annotation of transcript sequences is essential for transcript quantification and differential gene and transcript expression analysis. Single molecule long read sequencing technologies provide improved integrity of transcript structures including alternative splicing, and transcription start and polyadenylation sites. However, accuracy is significantly affected by sequencing errors, mRNA degradation or incomplete cDNA synthesis.

**Results:** We present a new and comprehensive Arabidopsis thaliana Reference Transcript Dataset 3 (AtRTD3). AtRTD3 contains over 160k transcripts - twice that of the best current Arabidopsis transcriptome and including over 1,500 novel genes. 79% of transcripts are from Iso-seq with accurately defined splice junctions and transcription start and end sites. We developed novel methods to determine splice junctions and transcription start and end sites accurately. Mismatch profiles around splice junctions provided a powerful feature to distinguish correct splice junctions and remove false splice junctions. Stratified approaches identified high confidence transcription start/end sites and removed fragmentary transcripts due to degradation. AtRTD3 is a major improvement over existing transcriptomes as demonstrated by analysis of an Arabidopsis cold response RNA-seq time-series. AtRTD3 provided higher resolution of transcript expression profiling and identified cold- and light-induced differential transcription start and polyadenylation site usage.

**Conclusions:** AtRTD3 is the most comprehensive Arabidopsis transcriptome currently available. It improves the precision of differential gene and transcript expression, differential alternative splicing, and transcription start/end site usage from RNA-seq data. The novel methods for identifying accurate splice junctions and transcription start/end sites are widely applicable and will improve single molecule sequencing analysis from any species.

## Background

Accurate gene expression analysis at the transcript level is essential to understand all aspects of plant growth and development and their responses to abiotic and biotic stress. The magnitude and dynamics of transcriptional and post-transcriptional re-programming of the transcriptome provide insights into the cellular complexity of responses to external and internal cues. This complexity can now be readily explored using high throughput RNA-sequencing (RNA-seq) technologies and vastly improved analytical methods and software programs. The ability to quantify the expression levels of individual transcripts from Illumina short-read RNA-seq data was revolutionised by the development of rapid and accurate non-alignment programmes, Kallisto and Salmon [1, 2]. However, Kallisto and Salmon require a reference transcriptome for accurate transcript quantification and the power of such analyses greatly depends on the quality and comprehensiveness of the reference transcriptome being used.

RNA sequencing using long read single molecule sequencing technologies, namely Pacific Biosciences (PacBio) and Oxford Nanopore sequencing, offers improved integrity of transcript structures. Single molecule sequencing has the advantage of being able to identify transcription start and end (polyadenylation) sites (TSS and TES, respectively), alternative splicing (AS), alternative polyadenylation (APA) and the correct combinations of different TSS, TES and splice junctions (SJs). However, sequencing errors are common in single molecule sequencing and mis-mapping of reads to the genome significantly increases false splice sites and affect open reading frames of transcripts [3]. Previous work on sequence alignment accuracy found that the main source of error for global sequence alignment was the misplacement of gaps, a phenomenon also called “edge wander” [4]. Misplacement of gaps is strongly affected by sequencing errors. Introns can be considered as “gaps” when the single molecule long reads are mapped to the genome and can generate many false splice junctions [5–9]. For example, alignment of high error-containing long reads from a particular locus often disagrees with one another (particularly around splice sites) [6] and high error rates result in a high proportion (27%) of mis-placed splice junctions [5]. Strategies to overcome the effects of sequencing errors in the long reads include by self- or hybrid correction methods. Self-correction utilizes the raw signal and consensus-based calls to reduce errors while hybrid correction exploits Illumina short reads to correct errors in the long reads [10–13]. However, current error correction tools tend to trim or split long reads when lacking local short read support, over-correct (introduce new, false splice junctions) when mapping to the wrong locations and lose isoforms with low expression [5, 7]. In addition, a considerable number of reads representing fragments of mature mRNAs, likely due to incomplete cDNA synthesis or mRNA degradation, compromise the accurate determination of transcription start and end (poly(A)) sites. While these issues are not generally appreciated, they reduce the overall precision of transcript quantification and downstream analysis of differential expression, AS, APA and TSS and TES usage.

Iso-seq single molecule sequencing has been applied to a wide selection of crop plants (e.g. maize, wheat, sorghum, coffee, tea, sugarcane, rice, amaranth [14] and grape), economically important plants for feed or products (e.g. switchgrass, Bermuda grass, perennial ryegrass, pine, rubber, red clover), wild plant species (e.g. wild strawberry), plants of botanical interest (e.g. Pitcher plants – Nepenthes spp.), and medicinal plants (e.g. Zanthoxylum, safflower, Salvia) [15,16,25–34,17,35–38,18–24]. The majority of these applications of PacBio sequencing investigated transcriptome diversity and complexity and determined transcription start sites, AS events and APA sites. However, significant issues surrounding the accuracy of SJs, TSS and TES identification suggest that the majority of the above transcriptome studies would benefit from improved methods of transcript structure determination. Accurate and well-curated transcripts also play an important role in improving genome annotations and the identification of novel genes and, particularly, long non-coding RNAs.

In this paper, we report the construction of a new, comprehensive Arabidopsis transcriptome, AtRTD3, based on a wide range of Arabidopsis tissues and treatments. AtRTD3 contains over 160k transcripts, 79% of which are derived from Iso-seq and have accurately defined SJs, TSS and TES. It improves the precision of analysis of RNA-seq data for differential gene and transcript expression and differential alternative splicing and now allows analysis of differential TSS and TES usage. We used a new pipeline based on TAMA [7] to analyse the Iso-seq data and developed novel methods to address the impact of sequencing errors and incomplete transcripts. We developed 1) a splice junction-centric approach that allows the identification of high confidence SJs and 2) a probabilistic 5’ and 3’ end determination method that effectively removes transcript fragments and identifies dominant transcript start and end sites. They allow accurate determination of SJs, TSS and TES directly from the Iso-seq data and remove the requirement for hybrid error correction or parallel experimental approaches for detecting TSS and TES such as CAGE-seq or poly(A)-seq, respectively. The defined sets of high confidence SJs, TSS and TES were used to generate an Iso-seq based transcriptome (AtIso) consisting of transcripts with accurately defined 5’ and 3’ ends and SJs and the combination of AS events with specific TSS and TES. The high confidence full-length transcripts in AtIso covered ca. two-thirds of genes in Arabidopsis and confirmed many of the short read assembled transcripts while resolving assembly artefacts present in AtRTD2 [39]. Around one-third of genes had very low or no Iso-seq coverage. Short read assembly generates highly accurate SJs but little information on 5’ and 3’ ends. Therefore, AtIso was merged with short read assemblies, such as AtRTD2 [39] and Araport11 [40] to form AtRTD3, giving preference to Iso-seq transcripts to capture high confidence SJs, TSSs and TESs and integrating only those transcripts from AtRTD2 and Araport11 with novel SJs or loci. The resulting AtRTD3 transcriptome contains 40,932 genes and 169,503 transcripts with ca. 78% of transcripts having Iso-seq support. AtRTD3 represents a significant improvement over existing Arabidopsis transcriptomes as demonstrated by its improved transcript quantification accuracy and transcript expression profiling over AtRTD2 and Araport11 and identification of differential TSS and TES usage induced by cold and light.

## Results

### Single molecule Iso-sequencing of diverse Arabidopsis plant samples

PacBio Iso-seq was performed on total RNA extracted from nineteen samples from different Arabidopsis Col-0 organs, developmental stages, abiotic stress conditions, infection with different pathogens and RNA degradation mutants to capture a broad diversity of transcripts (Additional File 1: Table S1). PacBio non-size selected Iso-seq libraries were made for all nineteen samples using a cap enrichment protocol (Teloprime, Lexogen). In addition,Teloprime v2 (Lexogen) libraries were constructed for six of the above RNA samples and Clontech (Takara Bio) libraries for two of the above samples. Each of the 27 libraries were sequenced on a separate SMRT cell on a PacBio Sequel machine with a 10 h (v3) movie time. The PacBio raw reads were processed using the PacBio IsoSeq3 pipeline to generate circular consensus sequences (CCS) and full length non-chimeric (FLNC) reads without the clustering and polishing steps and FLNCs were mapped to the reference genome (TAIR10) (Fig. 1). The numbers of reads,, FLNCs and mapped FLNCs along with statistics are shown in Additional File 1: Table S2. The 27 libraries generated 13.7 million Iso-seq reads in total. The total number of CCS was 8.7 M with an average of 322K CCS per library. The total number of FLNCs generated using lima+refine (see Methods) was 7.77 M with an average of 288K per library. About 7.36 M of the FLNCs mapped onto the Arabidopsis genome, generating 142.9K transcripts and 14.3K genes on average per library. We then merged the transcripts generated from the 27 libraries using TAMA merge, where unique transcripts including those with only a single nucleotide difference at 5’ and 3’ UTR were kept (Fig. 1). The merged transcriptome assembly consisted of 33,154 genes and 2,239,270 transcripts.

**Figure 1.**
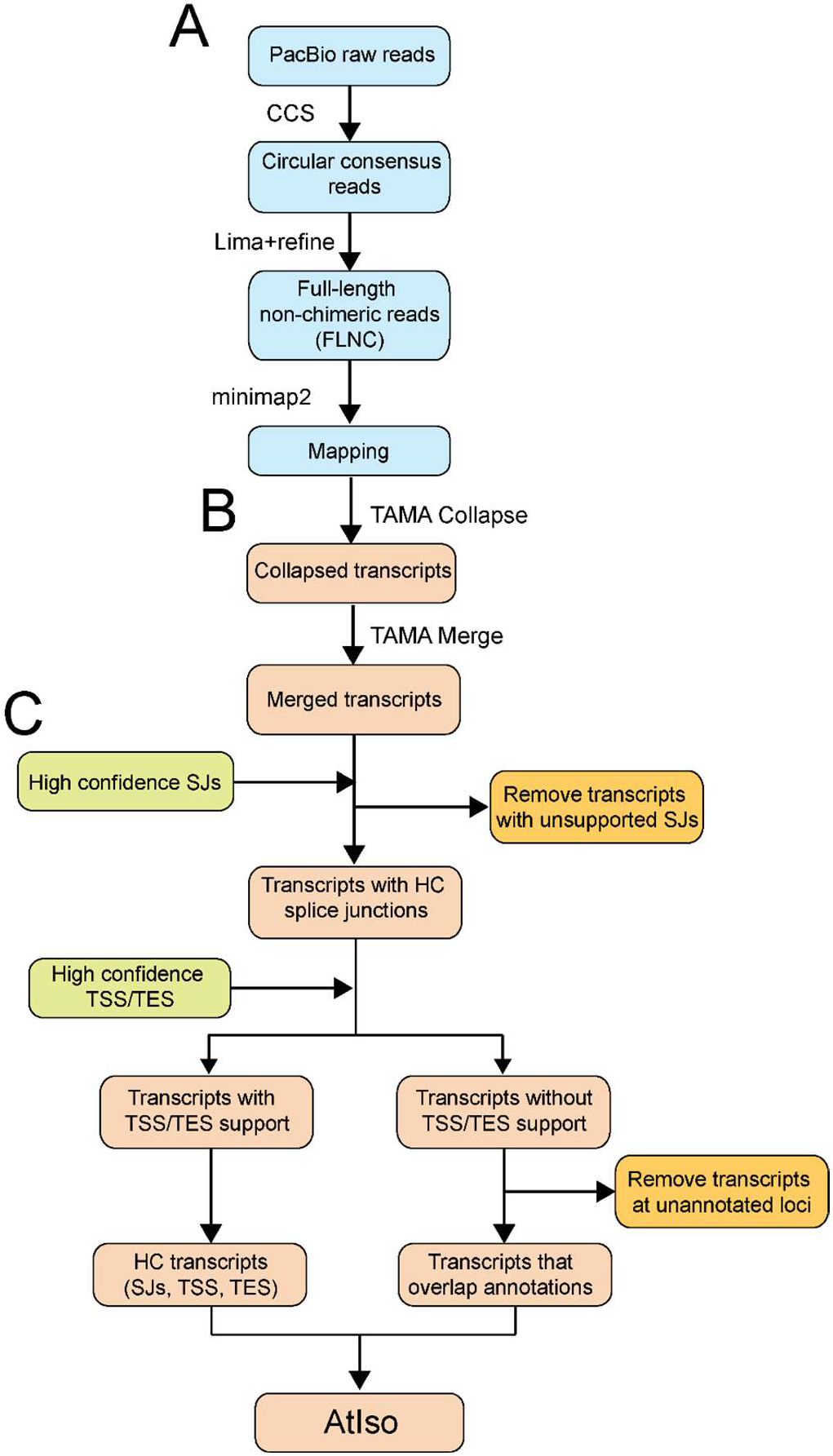
Workflow of analysis of PacBio Iso-sequencing. A) Raw reads are analysed using the PacBio Iso-seq 3 pipeline to generate FLNCs which are mapped to the genome (blue boxes). B) Mapped FLNCs are collapsed and merged using TAMA to generate transcripts (pink boxes). C) Transcripts are quality controlled using datasets of high confidence (HC) splice junctions (SJs) and transcript start and end sites (TSS/TES). Transcripts with unsupported splice junctions where reads contain errors within ±10 nt of an SJ are removed. Transcripts with both high confidence TSS and TES (determined by binomial probability for highly expressed genes and by end support with >2 reads for low expressed genes) are retained as HC transcripts. The remaining transcripts which have partial or no TSS and/or TES support were removed unless they overlapped with annotated gene loci. These transcripts, from genes with low expression and no/low coverage by Iso-seq, were combined with the HC transcripts to form AtIso (Arabidopsis Iso-seq based transcriptome).

### Sequencing mismatches around splice junctions (SJs) distinguish high and low confidence SJs

The challenge with Iso-seq derived transcripts is to accurately define SJs, TSSs and TESs. As the merged assembly contained tens of thousands of false SJs (see below), transcripts containing these SJs were identified and removed before defining TSS and TES. Based on the hypothesis that sequence errors in the Iso-seq reads around SJs promote “edge wander” [4] resulting in false SJs, we used TAMA Collapse to extract the mapping information of 30nt up and down streams of each SJs from the uncorrected reads from the 27 Iso-seq libraries. (Additional File 2: Fig. S1). We compared the resulting Iso-seq SJs to those of AtRTD2. 124,328 SJs were shared between Iso-seq and AtRTD2 transcripts and 110,992 were unique to the Iso-seq transcripts (Fig. 2A). We then extracted the mismatch profiles for the shared SJs and for those unique to Iso-seq transcripts and determined the number and percentage of mismatches in each position in the 30 nt up- and downstream of the SJ (Additional File 1: Tables S3A and S3B). Thus, the SJs in the Iso-seq transcripts were divided into two sets: 1) a high confidence set of 124,328 SJs (above) that were also present in AtRTD2 transcripts (extensive quality control measures were used to remove false SJs during the construction of AtRTD2 from short reads – Zhang et al. [39]) and 2) a low confidence set of 110,992 SJs unique to Iso-seq transcripts (above) that includes novel, bona fide junctions as well as incorrect mis-placed SJs. To assess the different characteristics between the two sets of SJs, we calculated position weight matrix (PWM) scores for 5’ and 3’ splice site consensus sequences for each intron (Additional File 1: Table S4). The average PWM scores of the high confidence SJs (5’ splice site: 69.91, 3’ splice site: 67.75) were significantly higher than the average of the low confidence set (5’: 62.79; 3’: 62.67) (Fig. 2B). Taking the threshold PWM of 65 as the criteria for a good quality splice site [39], 79.4% of high confidence SJs had PWM scores at both 5’ and 3’ splice sites of ≥65 with only 20.6% having at least one PWM score lower than the threshold (5’: 3.50% and 3’: 17.64%). In contrast, 79.17% of the SJs in the low confidence set have at least one PWM score lower than the threshold at either the 5’ or 3’ splice site (5’: 52.24% and 3’: 59.07%). Thus, the high confidence SJs have higher splice site consensus sequence quality characteristics than the low confidence SJs.

**Figure 2.**
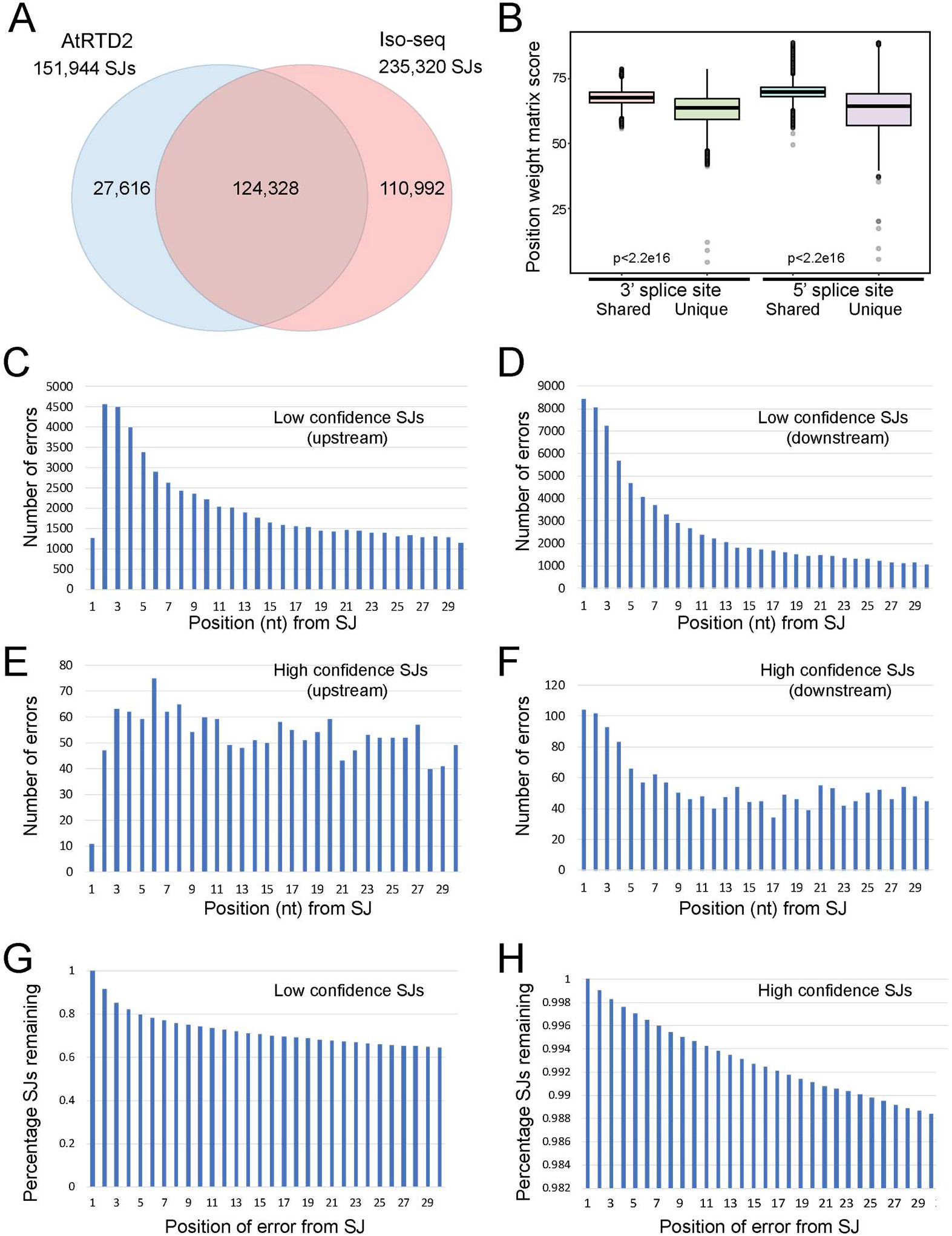
Impact of errors around splice junctions on the accuracy of their determination. A) Splice junctions (SJs) shared by AtRTD2 and Iso-seq (LDE_30; sjt_30) and unique to each. B) Position Weight Matrix (PWM) scores for splice sites unique to Iso-seq transcripts and shared with AtRTD2. PWM scores for 5’ and 3’ splice site sequences from SJs shared between AtRTD2 and Iso-seq transcripts (high confidence), are significantly higher (t- test, p < 2.26e-16) than those unique to Iso-seq (low confidence). C-F) Distribution of the number of errors in each position B,D) upstream and C,E) downstream of SJs unique to Iso- seq (low confidence) (B,C) and shared with AtRTD2 (high confidence) (D,E). G,H) Filtering of SJs based on errors in each position flanking SJs. The number of SJs remaining (as a percentage) after removal of SJs with errors in each position. F) Splice junctions unique to Iso-seq (low confidence) and G) SJs common to AtRTD2 (high confidence).

To examine the relationship between the presence of sequencing errors in reads around the SJs and the quality of the SJs, we selected the Iso-seq read with the smallest number of mismatches in the 60 nt region around each SJ for the analysis. The mismatch rate in each position in the SJs shared with AtRTD2 (high confidence set) was in the range of 0.008% to 0.08%. In contrast, the mismatch rate in each position in the low confidence SJs unique to Iso-seq were up to 100-fold greater and ranged from 1.02% to 4.12% (sequence upstream of SJ) and 0.97% to 7.58% (downstream of SJ) in, for the most part, descending order with distance from the splice junction (Additional File 1: Table S3A and S3B). Plotting the distributions of the mismatches at each position clearly showed a high number of mismatches in the vicinity of SJs unique to Iso-seq (low confidence) (Fig. 2C and D; Table S3A and B). Fewer mismatches with a more uniform distribution were observed for the SJs shared with AtRTD2 (high confidence) (Fig. 2E and F; Table S3A and B).

The effect of having sequencing errors in Iso-seq reads in the region of a SJ is illustrated by the number of SJs that would remain (recall) if SJs with an error in any of the positions were removed. For example, removing those SJs with a mismatch in positions 1-10 on either side of a SJ would remove only 711 SJs from the shared SJs (high confidence) leaving 99.43% of SJs (Fig. 2H; Additional File 1: Table S5A) but 29,606 SJs of the SJs unique to Iso-seq (low confidence), leaving 73.32% of the SJs (Fig. 2G; Additional File 1: Table S5B). Thus, sequencing mismatches in the vicinity of SJs are strongly associated with new, false SJs which carry over into transcripts. Filtering the SJs by removing those with mismatches around the SJs has a significant impact on the low confidence SJs but a very limited effect on the high confidence SJs. Thus, examining mismatches around the SJs is an effective strategy to distinguish high and low quality SJs and identify false SJs.

### Splice junction centric analysis for accurate splice junction determination

To apply the above observations to overcome the problem of false splice junctions being generated due to mismatches in Iso-seq reads in the vicinity of SJs, we developed a method to identify and retain high confidence SJs. The original TAMA collapse [7] removes reads with defined mismatches around the SJs. There are two issues with this approach: 1) when an Iso-seq read with multiple SJs is removed due to erroneous mapping of one or more SJs, other correct SJs supported by that read will be discarded at the same time; and 2) as Iso-seq sequencing errors are distributed randomly, some reads with errors around SJs could still be correct and be rescued by other reads that mapped perfectly to the region. We therefore modified the approach to keep all high confidence SJs irrespective of whether low quality SJs were present in the rest of the read. In so doing, we constructed a high confidence set of SJs where each SJ has support from at least one Iso-seq read with zero mismatches in positions ±10 nt from SJs. Using this set of SJs, reads with correctly mapped SJs but mismatches around SJs are still retained, contributing to identification of SJs in the final merged transcript assemblies.

In the transcript set from the 27 libraries, there were 235,320 non-redundant SJs. We first removed SJs with non-canonical motifs leaving 175,827 SJs. Then, we selected the SJs that had support from at least one read with zero mismatches to the genome in the 10 nt region on each side of the SJ. This reduced the number of false SJs caused by the combined effects of mis-mapping of the introns and sequencing errors around SJs, leaving 162,888 SJs. Thus, 71,726 (64.62%) SJs unique to Iso-seq (30.5% of all SJs in Iso-seq) were removed due to lack of experimental evidence for a high confidence SJ. For comparison, only 706 SJs that are shared between Iso-seq and AtRTD2 (0.46% of all SJs in AtRTD2) were removed using the same filtering parameters. Thus, the SJ-centric approach makes the best use of local information around the SJs of long reads to define the set of high confidence SJs.

### A stratified and probabilistic approach to determine the TSS and TES sites

The combination of Teloprime 5’ capture followed by Iso-seq sequencing from poly(A) tails should, in principle, produce full-length mRNA sequences containing authentic 5’-end/TSS and 3’-end/TES. However, a number of factors affect accurate TSS and TES identification: 1) mRNAs undergo degradation (in vivo or during RNA manipulation) generating truncated transcript fragments. Teloprime 5’ capture is not 100% efficient such that Iso-seq reads from 5’-degraded transcripts are still generated. Similarly, 3’ end degradation and off priming, where the PCR oligo-dT primer amplifies from poly(A) sequences within the transcript instead of the poly(A) tail, generate 3’ truncated transcript fragments. Thus, reads from transcripts with different degrees of degradation generate multiple false TSSs/TESs; 2) TSS/TES are usually stochastic and not limited to a single nucleotide location but rather are distributed around a dominant site [41]; and 3) the number of Iso-seq reads varies greatly across a large dynamic range. Consequently, highly expressed genes may contain thousands of individual transcripts including substantial numbers of degradation products. In contrast, for genes with low levels of expression and a limited number of reads or no read coverage, it is difficult to apply statistical inference to determine whether read start/end points are TSSs/TESs. The challenges in accurately identifying TSS/TES for genes with high and low read abundance are therefore very different. For highly sequenced genes, the major task is to reduce false TSS/TES from transcript fragments and identify dominant sites. For genes with few reads, the task is to get sufficient experimental evidence to support TSS/TES identification. We have therefore developed and applied two different approaches to end determination depending on the read/transcript abundance. We assumed that for highly squenced genes, authentic TSS/TES sites would tend to be sequenced more often while the ends from degraded mRNA products would occur randomly. We, therefore, used the binomial function to estimate the probability of having a certain number of Iso-seq read ends at any position at random and used these probabilities to identify positions with non-random (i.e. enriched) ends that represent authentic start or end sites (Additional File 2: Fig. S2A and B). For genes with few reads, we compared start and end sites of different reads to identify similar ends as support for potential TSS/TES (see below; Additional File 2: Fig. S2C).

### Identification of significant read start and end genomic locations

For TSS determination, only the Teloprime captured reads were used as the Clontech libraries are more likely to contain truncated fragments with missing TSSs [42]. The exact genomic co- ordinate of the start of each read (read start genome location-RSGL) was identified giving a total of a total of 616,593 RSGLs. By applying the binomial probability method (Additional File 2: Fig. S2A), 61,014 significant RSGLs enriched for start locations were detected from 17,098 genes. These 17,098 genes tended to be highly sequenced with read numbers ranging from 7 to 48,110 and a median of 110 reads. They accounted for 550,022 of the total RSGLs (89.3%) from which the 61,014 significant non-random RSGLs were identified, an approximately 10-fold reduction of the average RSGL number per gene from 32.17 to 3.57. Thus, the binomial probability method reduces the overall number of RSGLs into a smaller number of high confidence RSGLs. For the remaining 15,858 genes, no significant RSGLs were detected. These genes had relatively few reads with a median of 2 reads per gene and 80% of genes having fewer than 7 reads. For these genes, we compared the start positions of reads from each gene and required at least two Iso-seq reads with 5’ ends within a sliding window of 11 nt (5 nt on each side) to call a supported RSGL (Additional File 2: Fig. S2C). By this method, the 66,571 remaining RSGLs (from 15,858 low abundance genes) generated 25,930 supported RSGLs from 7,028 genes. Thus, we have defined 61,014 and 25,930 TSSs from with high and low numbers of reads genes, respectively.

Before enrichment, a total of 723,903 read end genomic locations (REGLs) were identified. We removed 11,703 reads where 3’ ends were immediately followed by poly(A) sequences in the genome sequence and were likely to be a result of off-priming, leaving a total of 712,200 REGLs. We then applied the binomial distribution method to detect non-random REGLs, as described above for RSGLs. For highly sequenced genes, 84,043 significant REGLs (Additional File 2: Fig. S2B) from 16,728 genes were identified with read abundance per gene varying from 7 to 49,917 and a median of 128. These highly sequenced genes contained 669,642 (94.02%) of the total REGLs and showed very variable end sites. The binomial distribution probability method reduced the average number of REGLs per gene from 40.03 to 5.02. The remaining 13,440 genes had fewer reads with a median of 1 read, and 80% of these genes had fewer than 5 reads. At least two Iso-seq long reads with similar 3’-ends within a sliding window of 11 nt (5 nt on each side) were required to call a supported REGL (Additional File 2: Fig. S2C). On this basis, from the 42,558 REGLs from the 13,440 genes with few reads, 21,664 supported REGLs from 5,824 genes were identified. Thus, we have defined 84,043 significant REGLs and 21,664 supported REGLs from genes with high and low numbers of reads, respectively. Finally, 8,830 and 7,616 genes did not have significant or supported RSGLs or REGLs, respectively, and represented genes with one read or with very few reads with varying start or end locations differing by more than 5 nt.

### Validation of significant TSSs and TESs

A transcription start site dataset for Arabidopsis genes at nucleotide resolution was generated previously using paired-end analysis of TSSs (PEAT) [41]. In their study, using a pooled Col- 0 root sample, 79,706 mapped and annotated PEAT tag clusters (groups of peaks) were identified and quality filtering generated 9,326 strong tag clusters from protein-coding genes where TSS locations were supported by at least 100 reads. The information for each tag cluster included the start, end, strand, as well as the mode, which is the location within the cluster where the greatest number of 5’ ends were mapped [41].

We compared the significant 61,014 RSGLs from the highly sequenced genes with the PEAT tag clusters and found that 43,254 (70.9%) were located within 14,957 of the raw PEAT clusters and 30,981 within 8,445 (90.5%) of the 9,326 strong tag clusters (Additional File 2: Fig. S3A, B). Thus, the significant RSGLs from the highly sequenced genes showed substantial concentration and overlap with the published set of strong tag clusters. We also compared the significant RSGLs with the mode (genomic locations with the highest number of reads within that tag cluster) of strong tag clusters and found that 6,563 (70.4%) strong tag cluster modes co-located with the significant RSGLs with no more than a 1 nt difference (Additional File 2: Fig. S3C). Significant RSGLs were also identified in another 9,010 genes not detected in Morton et al. [41]. This is likely due to the much wider range of tissues and treatments used here and differences in gene coverage between the Iso-seq and PEAT analyses. We have also compared our RSGLs to a recent study [43] that carried out genome-wide TSS mapping using 5’ CAP sequencing. In this study, 96,232 TSS tag clusters were detected in 21,359 genes in wild type plants and mutant lines of the FACT (FAcilitates Chromatin Transcription) complex. We found that 55,737 (91.35%) of our significant RSGLs located within the TSS tag clusters of Nielsen et al. [43], covering 16,353 genes. The correspondence of our data with both the above studies shows the high accuracy of the RSGLs detected using our novel method of transcript 5’ end determination.

Arabidopsis polyadenylation sites have been previously identified through direct RNA sequencing of seedling RNA, which found 49,916 cleavage and polyadenylation site (CS) peaks supported by >9 raw reads from 14,311 genes [44]. We compared the 84,043 significant REGLs with the CS peaks and found that 1) 45,931 (92%) CS peaks from 13,443 genes co-located with significant REGLs within a 50 nt window and 24,927 (49.93%) CS peaks co-located with significant REGLs at the same genomic location (≤ 1 nt difference) (Additional File 2: Fig. S3D, E). The significant REGLs identified an additional 12,305 TES sites in 5,531 genes, including 3,663 genes for which no CS peaks were reported [44]. The increased diversity of TES identified from our Iso-seq data are again likely due to the wider range of tissues and treatments used for RNA sequencing.

### Significant RSGLs/REGLs show enrichment in motifs related to TSSs and TESs

To further validate the TSSs in the significant 61,014 RSGLs, we looked for common transcription motifs (e.g. TATA box, lnitiator and Y-patch) in the region of the TSSs (+500 to - 500 bp) and the Kozak translation start site motif downstream of the TSS, and compared these to the raw 79,706 TSS tag cluster peaks from Morton et al. [41]. The TATA box is a T/A-rich motif ca. 25-35 bp upstream of highly expressed genes that determine expression levels [41,45,46] (Additional File 1: Table S6). A sharp peak was observed upstream of the TSSs for both RSGLs and TSS peak clusters consistent with the expected position for a TATA box (Fig. 3A). Thus, there is a good corroboration of our computational derived TSS and the experimentally defined TSS. Despite fewer significant RSGL sites being investigated, we found the number of TSSs with upstream TATA motifs in the RSGLs almost doubled that seen with the PEAT tag cluster peaks (from 3,603 sites to 6,976 sites) (Fig. 3A). A proportion test shows that the TATA box motif was significantly enriched in RSGLs compared to the TSS cluster peaks in Morton et al (p<2.2e-16). The Initiator (Inr) element is pyrimidine-rich, overlaps the TSS site, and is important for transcriptional activation [46] while the Y-patch pyrimidine-rich motif, found upstream of TSSs, is unique to plants and found in more than 50% of annotated rice genes [47] (Additional File 1: Table S6). Enrichment of both motifs around the TSS was observed again, with 2,236 and 11,477 instances, respectively, in the significant RSGL set and 1,208 and 6,067 instances, respectively, in the Morton et al. data (both proportion tests p<2.2e-16) (Fig. 3B and C). Finally, the Kozak consensus translation start sequence is downstream of the TSS and contains the translation start AUG codon [48]. Significant enrichment of Kozak sequences was seen downstream of the TSSs for the significant RSGLs with 316 instances over 116 instances in the Morton data (proportion tests p<2.2e-16) (Fig. 3D).

**Figure 3:**
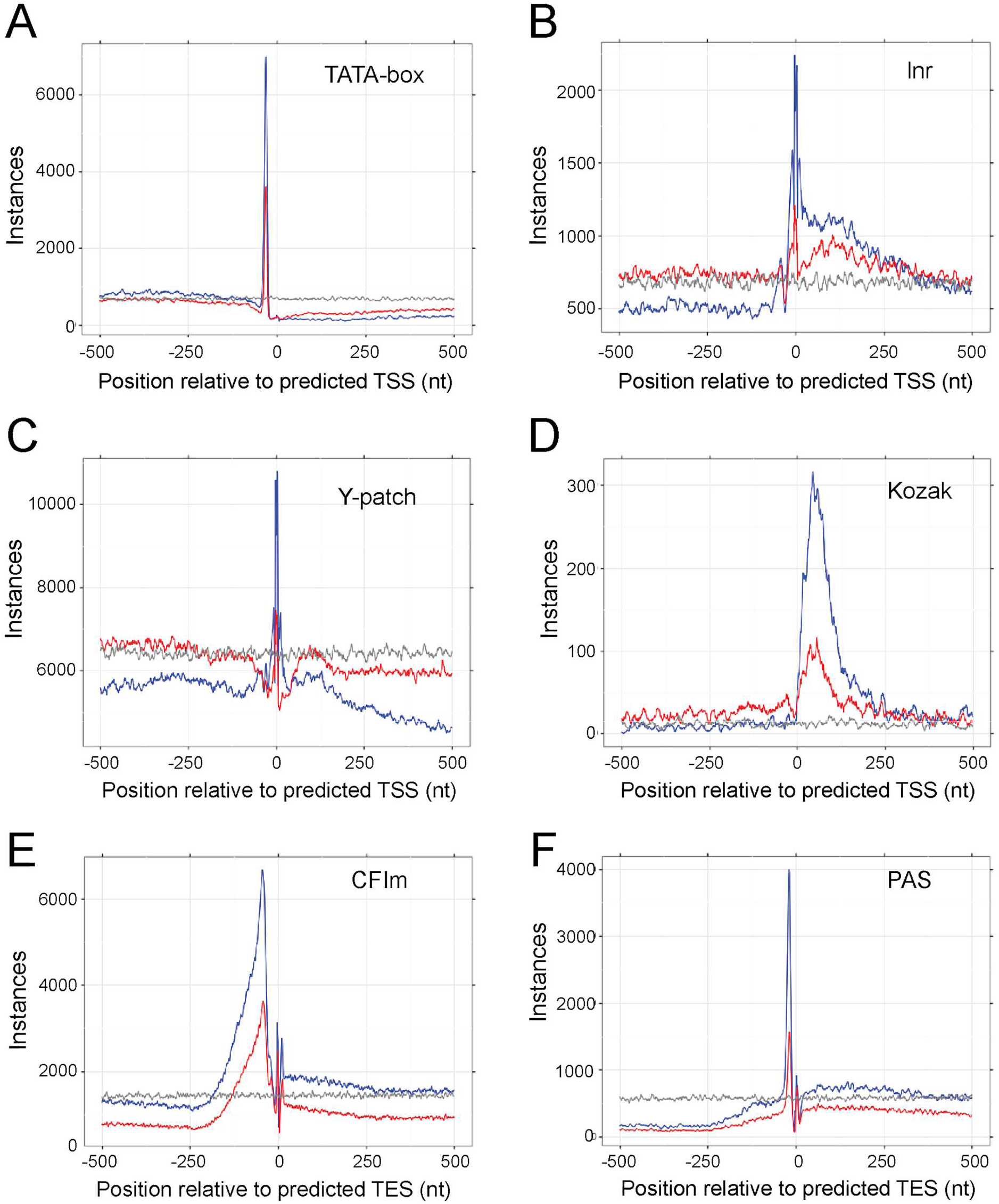
Enrichment of sequence motifs associated with TSS and TES sites. A-D) TSS sites: A) TATA box, B) Initiator (Inr), C) Y-patch, D) Kozak translation start site consensus motif; E-F) TES sites: E) CFlm and F) PAS. Lines indicate instances of motif in relation to start and end sites from Iso-seq (blue), Morton et al. (2014) A-D, and Sherstnev et al. (2012) E,F (red); random control (grey).

To further validate the REGLs, we searched the genomic sequences around the TESs (-500 to +500 bp) of the 84,043 significant REGLs and 49,916 CS peaks from Sherstnev et al. [44] for conserved cleavage and polyadenylation sequence motifs. The polyadenylation signal (PAS) motif (possessing a canonical AATAAA when the PAS is relatively strong) is required for 3’ end polyadenylation while the CFlm motif is the binding site of cleavage factor Im, an essential 3’ processing factor [49, 50] (Additional File 1: Table S6). The number of matching sequences and their positions showed significant enrichment of CFIm sequences (upstream of the PAS) and the poly(A) signal motif at the TES for significant REGLs over CS peaks (Fig. 3E and F, respectively) with 3,627 and 1,565 instances, respectively, in the CS peaks and 6,663 and 3,994 instances, respectively, in the significant REGLs (proportion tests p = 5.725e- 06 and p<2.2e-16, respectively) (Fig. 3E and F).

### Generation of high level transcripts for AtIso

To achieve accurate transcript isoforms from the Iso-seq data, we have adopted a strategy that requires evidential support for all the SJs, TSSs and TESs. We generated datasets of high confidence SJs, RSGLs and REGLs which were then used to filter the 2,239,270 transcripts from all the libraries. Given the stochastic nature of TSSs and TESs, we applied a 100 nt window around each significant and supported RSGL and REGL (50 nt on each side) to define high confidence TSS and TES regions (Additional File 2; Figure S4). This generated 1,674,795 transcripts after sequentially removing transcripts containing poorly supported SJs (117,361 transcripts) or poorly supported TSS and TES (447,114 transcripts) (Fig. 1C). The above filtering criteria also addressed the common issue of excessive numbers of single exon gene models generated from Iso-seq experiments and many other genome-wide annotation projects [7, 51], which could be the result of genomic DNA contamination. In our data, we also observed that 161,578 (46.6%) out of 346,455 single exon transcripts were removed due to the TSS/TES filtering. These removed transcripts are probably fragments with missing 5’ or 3’ sites or false positive gene models. As a result, filtering using high confidence TSS and TES regions also reduced the number of the mono-exonic genes (containing mono-exonic transcripts) from 13,619 to 4,477, a reduction of 67% on the number of putative mono-exonic genes. The percentage of mono-exonic genes decreased from 41.3% to 20.9% of the total number of genes after TSS/TES filtering.

Finally, to increase the gene coverage using existing annotations and make the maximum use of the Iso-seq long reads, we retained a further 2,483 genes (7,398 transcripts) where the reads covered Araport transcripts on the same strand with at least 50% overlap. The combined set was merged allowing 50 nt variations at the 5’ and 3’ ends and the final AtIso dataset contained 24,344 genes with 132,190 high level transcripts (Additional File 2: Fig. S4).

### Construction and characterisation of the AtRTD3 transcriptome

AtIso contained transcripts from 57% of the genes in Araport11. Splice junction and transcript identity were compared among AtIso, AtRTD2 and Araport11 [39, 40] (Additional File 3). There was high similarity in SJs but very low overlap of transcripts due to poor 5’ and 3’ end determination and different combination of SJs in AtRTD2 and Araport11 compared to the Iso-seq transcripts (Additional File 3). To generate a new, comprehensive transcriptome for Arabidopsis that covered all genes and incorporated the Iso-seq transcripts, long and short read assemblies were combined using the following criteria: 1) AtIso had the most accurate transcript data and was used as the back-bone for integrating AtRTD2 and Araport11. To maximize the use of Iso-seq transcripts, we kept all AtIso transcripts; 2) As the TSS, TES and the combination of SJs are less accurate in transcripts assembled from short reads, a) only transcripts from AtRTD2 and Araport that contained novel SJs or b) covered novel genomic loci were incorporated from the short read assemblies. Using these criteria, the three assemblies were merged with TAMA merge, generating the final transcriptome, which we named AtRTD3. AtRTD3 contained 40,932 genes with 169,503 transcripts with a total of 183,568 SJs. AtIso contributed 132,166 (77.97%) transcripts from 25,248 (61.68%) genes, AtRTD2 contributed 24,831 (14.65%) transcripts from 13,683 genes [39] and Araport11 contributed 12,506 (7.38%) transcripts from 11,750 genes [40]. In AtRTD3, the average number of isoforms per gene was 4.4 and nearly 80% of transcripts had Iso-seq support (SJs, TSS and TES).

Genes and transcripts in AtRTD3 were characterised using TranSuite, a program which identifies mono- and multi-exonic genes and generates accurate translations of transcripts and transcript characteristics [52]. The output includes translations of all transcripts in the RTD and multiple transcript features (Additional File 1: Table S7). These results are summarised in Fig. 4 and Additional File 1: Table S8A and S8B. Almost three-quarters (73.5%) of the genes coded for proteins and ca. 26.5% were non-protein-coding genes (Fig. 4A; Additional File 1: Table S8A). Of all genes, 66.5% were multi-exonic and 50% had more than one transcript isoform. Of the genes that produced a single transcript, two-thirds were single exon genes and one third were multi-exonic (Fig. 4C; Additional File 1: Table S8B). For protein-coding genes, 62.9% were multi-exonic with more than one isoform. The 10,827 non-protein-coding genes generated 14,880 transcripts (Fig. 4E); the majority were single exon genes but 1,728 genes were multi-exonic (spliced) with a single transcript and over 5k genes had more than one isoform (Additional File 1: Table S8A). We also identified 3,796 chimeric (read-through) transcripts covering usually two Araport genes with an overlap > 30%.

**Figure 4.**
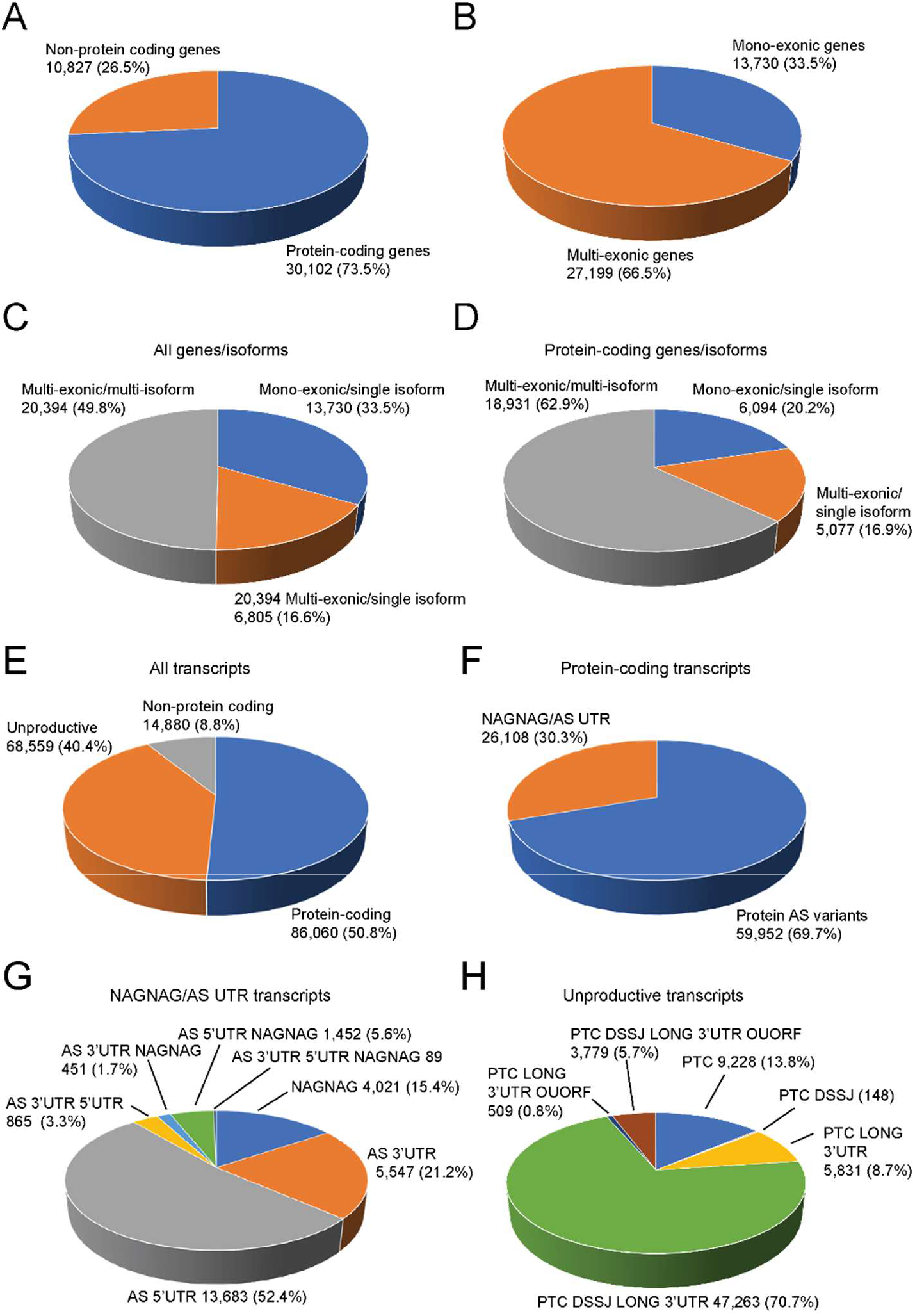
Gene and transcript characteristics of AtRTD3. A) Protein-coding and non-protein-coding genes; B) Mono-exonic and multi-exonic genes; C) Mono- and multi-exonic genes with single/multiple transcript isoforms for all genes and D) for protein-coding genes; E) distribution of transcripts from protein-coding genes (protein-coding and unproductive isoforms) and from non-protein-coding genes; F) Protein-coding transcripts with little or no impact on coding sequence (NAGNAG/AS in UTR) and protein-coding variants; G) distribution of transcripts with NAGNAG, AS in 5’ UTR and AS in 3’ UTR: H) distribution of NMD features among unproductive transcripts from protein-coding genes. DSSJ - downstream splice junction; OUORF - overlapping upstream open reading frame.

At the transcript level, AtRTD3 contained more than double the number of transcripts compared to AtRTD2 with greatly increased numbers of protein-coding and unproductive transcripts from protein-coding genes: 154,619 (91.2%) AtRTD3 transcripts came from protein-coding genes (Fig. 4E). Of these, ca. 86K are expected to code for proteins while ca. 68.5K are probably unproductive (Fig. 4E; Additional File 1: Table S8B). Alternatively spliced transcripts that coded for proteins were divided into those where AS events had little or no effect on the coding sequence (NAGNAG/AS UTR) (30.3%) and those that encoded protein variants (69.7%) (Fig. 4F; Additional File 1: Table S8B). NAGNAG/AS events generate transcripts that code for protein variants differing by only one amino acid and transcripts of genes where AS events occur only in the 5’ and/or 3’ UTRs and hence code for identical proteins. The NAGNAG/AS UTR transcripts were further broken down according to whether AS events were in the 5’ and/or 3’UTR or were NAGNAG (Fig. 4G; Additional File 1: Table S8B). The most frequent AS events were in the 5’ UTR (52.4%) followed by those in the 3’ UTR (21.2%) or NAGNAG events (15.4%) (Fig. 4G). NAGNAG AS events were present in 7% of protein-coding transcripts and 3.5% of all transcripts. Finally, the unproductive transcripts from protein-coding genes were classified by their nonsense mediated decay (NMD) target features: presence of a premature termination codon (PTC), downstream splice junctions, long 3’ UTR, or overlapping upstream ORF where an upstream ORF overlaps the authentic translation start site [53] (Fig. 4H; Additional File 1: Table S8B). Over 70% of the unproductive transcripts contained the classical combination of NMD target features of a PTC with downstream splice junctions and long 3’UTRs, 8.7% had a PTC with either one of these signals and 6.4% of transcripts contained an overlapping uORF (Fig. 4H; Additional File 1: Table S8B).

Iso-seq increased the number of transcript isoforms for many genes reflecting both discovery of novel AS events and defined TSS/TES variation compared to Araport and AtRTD2 (Additional File 2: Fig. S5). Different TSS in Iso-seq transcripts were observed in genes where alternative TSS had been previously characterised [54], for example, AT1G09140 (SERINE-ARGININE PROTEIN 30) and AT1G22630 (SSUH2-LIKE PROTEIN) (Additional File 2: Fig. S6A and B). Defined Iso-seq TESs in AtRTD3 confirmed the well-established intronic alternative polyadenylation sites in FCA and FPA (not shown) and those in ATHB13 (AT1G69780) and ANKYRIN REPEAT-CONTAINING PROTEIN 2 (AT4G35450) [55] (Additional File 2: Fig. S7A and B). The Iso-seq data also identified novel splice sites and alternative TSS/TES in known and novel lncRNAs. For example, AS transcripts of the antisense lncRNA, FLORE [56] were confirmed (Additional File 2: Fig. S8). AtRTD3 contained 1,541 novel genes compared to Araport (Additional file 1: Table 9A). All were identified by Iso-seq and their transcripts therefore have high confidence TSS/TES and SJs for those which are spliced or alternatively spliced. The majority of the novel genes were lncRNAs with only 109 genes coding for proteins with a CDS of >100 amino acids; 223 had more than one transcript and 1,318 had single transcripts. The novel genes were either intergenic or antisense genes. For example, G12636 is an alternatively spliced intergenic lncRNA, G13263 is a spliced antisense gene with different TSS and G14744 is an alternatively spliced antisense gene which covers two different protein-coding genes (Additional File 2: Fig. S9A, B and C, respectively). Finally, 1,197 genes had Iso-seq-defined 3,796 chimeric transcripts which extended over two or more genes (Additional file 1: Table 9B). For example, Iso-seq detected only a single transcript of the upstream MEKK2 gene but multiple chimeric transcripts covering the tandemly arranged MEKK2 and MEKK3 genes (Additional File 2: Fig. S10). Thus, the high quality Iso-seq data increases transcript diversity and provides detailed information of transcript features.

Finally, we compared the frequency of different AS event types among the different transcriptomes using SUPPA2 [57]. AtRTD3 had the highest number of AS events followed by AtIso (Additional File 1: Table S10). For the most part the frequency of different AS events is similar with approximately double the number of alternative 3’ splice site (Alt 3’ss) than alternative 5’ splice site (Alt 5’ss) events and relatively few exon skipping events (6-7%). Intron retention (IR) is far more frequent in plants than in animals with around 40% of plant AS events being IR [58] as seen in AtRTD2 and Araport11 (Additional File 1: Table S10). However, AtIso contained a higher number of IR events (50%) which supported the observation that many Iso-seq transcripts from multi-exon genes contained different individual retained introns (e.g. Additional File 1: Fig. S5 and S6) such that Iso-seq appeared to identify more low abundance transcript variants in highly expressed genes. Finally, the intermediate value of 44% IR events in AtRTD3 reflects the combination of unique transcripts from Iso-seq and short read-derived assemblies.

### AtRTD3 and AtIso increase quantification accuracy at the transcript and alternative splicing levels

To evaluate AtRTD3 and AtIso in the performance of transcript and AS quantification, we used high resolution (HR) RT-PCR data that we had used previously to evaluate AtRTD2 [39]. The HR RT-PCR data was generated using RNA samples of two time-points (T5 and T20) of Arabidopsis plants exposed to cold and which were also used to generate RNA-seq data for direct comparison (Calixto et al., 2018). Due to the increased transcript/AS diversity in AtRTD3 and AtIso, we were able to analyse 226 AS events from 71 Arabidopsis genes (three biological replicates of each of the T5 and T20 time-points). This generated 1,349 data points, which represents a significant increase from the earlier study (127 AS events from 62 genes with a total of 762 data points). For the splicing ratios from HR RT-PCR, transcript structures from AtRTD3 and AtIso were compared to the amplicons in HR RT-PCR and the TPMs of individual transcripts covering the different AS outcomes were used to calculate splicing ratios for each of the AS events or event combinations in that region. For splicing ratios from RNA-seq data, each of the different reference transcriptomes (AtRTD2-QUASI, Araport11, AtIso and AtRTD3) were used to quantify transcripts using Salmon. The splicing ratio for each AS event was calculated by comparing the abundance of individual AS transcripts with the AS event to the fully spliced (FS) transcript which is usually the most abundant transcript and codes for full-length protein (AS/FS). The scatter plot of splicing ratios from HR RT-PCR and RNA-seq using the different reference transcriptomes (Fig. 5) shows that AtRTD3 and AtIso achieve the highest concordance with HR RT-PCR data. This is likely due to the increased integrity of transcript structure (accurate characterization of SJs, TSSs and TESs and their combinations) as well as increased transcript/AS diversity over AtRTD2 and Araport11. Although AtIso and AtRTD3 performed very similarly in this analysis, AtRTD3 is the transcriptome of choice for RNA-seq analyses due to its far greater gene coverage.

**Figure 5.**
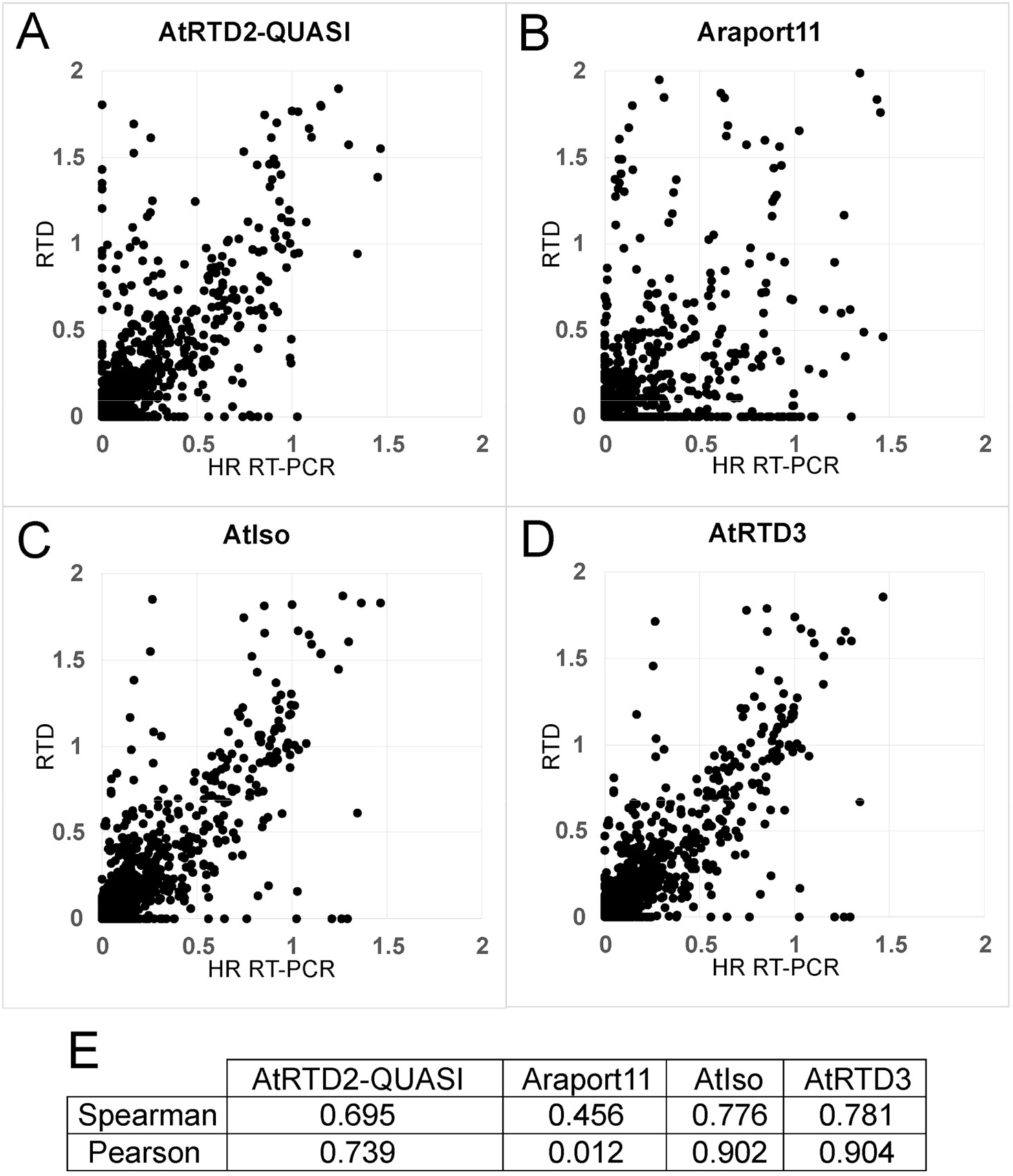
Correlation of splicing ratios calculated from the RNA-seq using different RTDs and HR RT-PCR data. Splicing ratios for 226 AS events from 71 Arabidopsis genes (three biological replicates of the time-points T5 and T20) generated 1349 data points in total. The splicing ratio of individual AS transcripts to the cognate fully spliced (FS) transcript was calculated from TPMs generated by Salmon and A) AtRTD2-QUASI, B) Araport11, C) AtIso and D) AtRTD3 and compared to the ratio from HR RT-PCR. E) Correlation coefficients are given for each plot. Note that for clarity of the figures, data-points with values that lie substantially outside the range of the graphs are not included in A-D) but are included in the correlation values.

### High resolution gene and transcript expression profiling with AtRTD3

AtRTD3 contains many more transcripts (169,503) than AtRTD2 (82,190). This reflects increased numbers of transcripts with intron retention and other AS events as well as defined TSS and TES variation. For some highly expressed genes with multiple introns, the combination of TSS/TES variation and intron retention events often led to tens of transcript isoforms from a single gene. Although more complex than AtRTD2, we predicted that the majority of isoforms with intron retention represent intermediates of splicing where an intron(s) had not been removed at the time of RNA extraction and that they would therefore have low levels of expression. Similarly, some isoforms with novel AS events would be NMD-sensitive again potentially with low expression levels. In contrast, novel AS isoforms or isoforms with different TSS or TES with significant expression levels would be expected to alter the transcript expression profiles compared to analysis with AtRTD2 where these isoforms were absent (we showed previously the impact of missing transcripts in transcript quantification - Zhang et al., 2017). To demonstrate the increased resolution obtained with the more complex and diverse AtRTD3, we compared gene and transcript expression profiles using RNA-seq data from an RNA-seq time-course of 5-week-old Arabidopsis plants grown in 12 h dark:12 h light in the transition from 20 ⁰C to 4 ⁰C [59, 60]. Briefly, transcripts were re-quantified with Salmon using AtRTD3 as reference and the RNA-seq data from 26 time-points (3 biological replicates) was re-analysed. Time-points were taken every 3 h for the last day at 20 ⁰C (T1-T9), the first day at 4 ⁰C (T10-T17) and the fourth day at 4 ⁰C (T18-T26) (see Fig. 6). Expression profiles were directly compared between AtRTD2 and AtRTD3.

**Figure 6.**
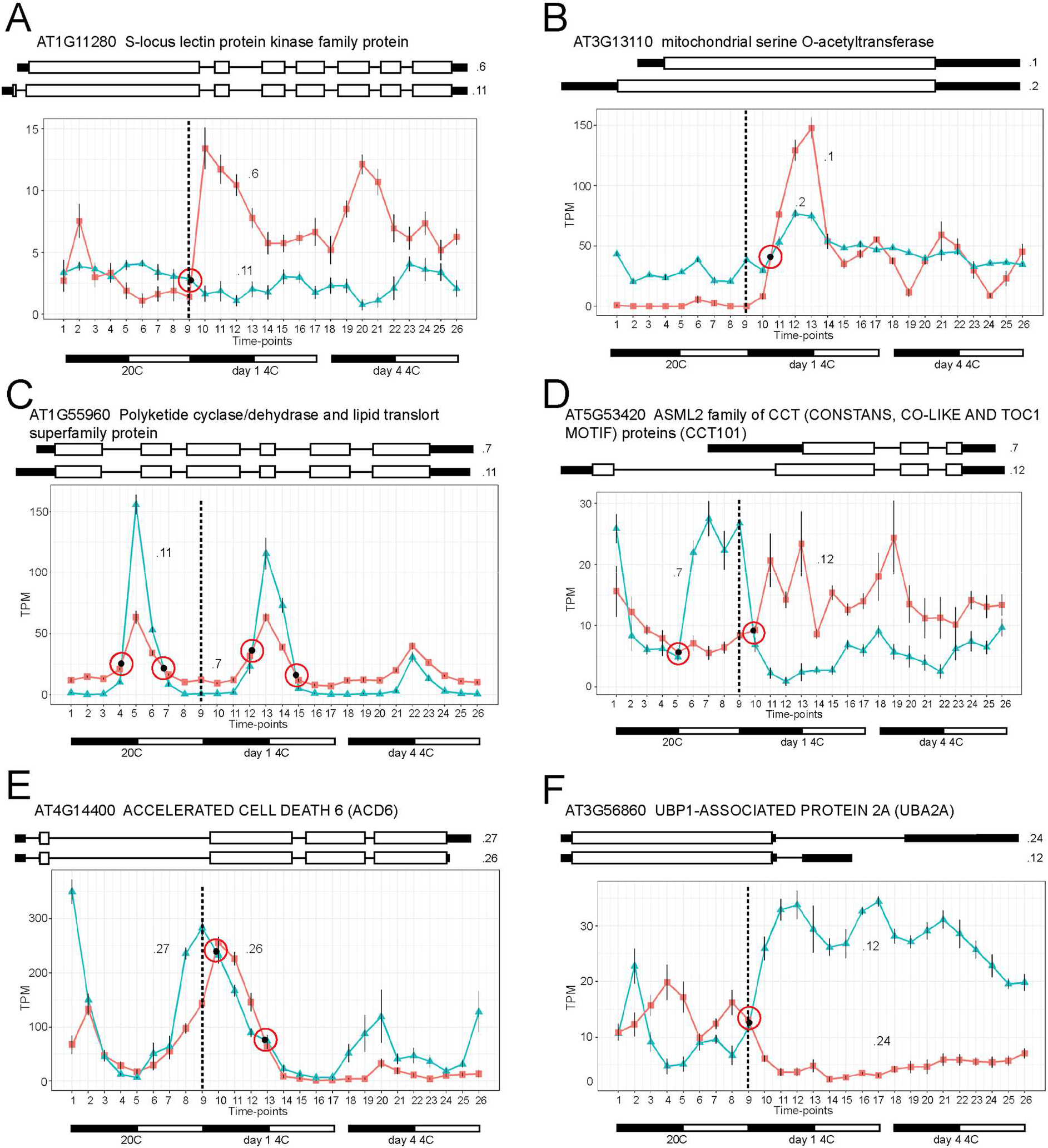
Differential TSS and TES usage. Pairs of transcript isoforms with significant isoform switches and different TSS (A-D) and TES (E and F). A) AT1G11280 – the shorter .6 transcript is cold-responsive; B) AT3G13110 – single exon gene with different TSS where the .1 transcript has rapid cold-induced expression compared to the .2 transcript; C) AT1G55960 – both transcripts peak at dusk but have different expression behaviour with the .11 isoform showing large increases of expression at 20 ⁰C and day 1 at 4 ⁰C declining with continued cold exposure; D) AT5G53420 – isoforms with very different TSS - .7 isoform expressed rhythmically peaking during the day (light-responsive) at 20 ⁰C before declining rapidly in the cold while the .12 transcript has increased expression in the cold, peaking during the dark; E) AT4G14400 – the isoforms differ only in their TES but are expressed rhythmically with different phase (3 h offset) at 20 ⁰C and reduced at 4 ⁰C; F) AT3G56860 – very different TES and expression behaviour – antiphasic at 20 ⁰C with cold-induced switch to the shorter .12 isoform. Error bars on points are standard errors of the mean.

The more comprehensive nature and accuracy of AtRTD3 is clearly illustrated by the THIAMIN C SYNTHASE (THIC) gene (AT2G29630) which is involved in regulation of thiamin biosynthesis via a riboswitch in the 3’ UTR that controls expression through alternative 3’-end processing or splicing [61, 62]. Three types of transcripts have been identified previously: Type I transcripts represent precursor transcripts; type II transcripts have been processed at a polyadenylation site in the second 3’UTR intron ((3’-2) and type III transcripts have splicing of intron 3’-2 (Wachter et al., 2007; Additional File 2: Fig. S11A). Low levels of THIC expression reduce vitamin B1 (thiamin diphosphate - TPP) levels. Low levels of TPP allow the structure of the RNA aptamer to interact with the 5’ splice site of the 3’-2 intron to inhibit splicing and promote processing at the polyadenylation site in the intron. The resultant type II RNA transcripts have relatively short 3’ UTRs, are stable and give high expression of THIC [61]( Additional File 2: Fig. S11A). With increased levels of TPP, TPP binds to the aptamer leading to structural changes in the riboswitch RNA such that it can no longer interact with and inhibit use of the 5’ splice site of 3’-2. Subsequent splicing of the 3’-2 intron removes the poly(A) site and type III transcripts with longer 3’ UTRs of various lengths are generated leading to increased RNA degradation and reduced expression of THIC (Additional File 2: Fig. S11A). AtRTD3 contained 32 THIC transcript isoforms (Additional File 2: Fig. S11B). The majority have very low expression and either have retention of different introns within the CDS and are likely intermediates of splicing or have other AS events that disrupt the ORF and introduce PTCs. Type I, II and III transcripts [61] were clearly distinguished by their 3’ UTR structures (Additional File 2: Fig. S11B). The 3’ processed type II mRNAs have a shorter 3’UTR than types I and !! due to processing at the pA site within intron 3’-2 while type III transcripts have splicing of the 3’-2 intron (removes the first seven nucleotides of the aptamer sequence) and longer 3’UTRs with a range of 3’ends sites [61]. In addition to the type I, II and II isoforms found in AtRTD3, we observed a novel AS variant where splicing removed only the first aptamer nucleotide. We detected three type I precursor transcript isoforms among the 32 THIC isoforms in AtRTD3 (Additional File 2: Fig. S11B). In contrast, Araport and AtRTD2 contained 4 and 10 transcripts, respectively. Neither AtRTD2 nor Araport contained type II transcripts and possible type I transcripts were much longer than those obtained with Iso-seq suggesting that the 3’UTRs of the transcripts were incorrectly assembled. THIC is highly expressed and under circadian control [62]. In the cold time-series analysed with AtRTD3 as reference, THIC expression increased during the day and decreased in the dark (Additional File 2: Fig. S11C). The major isoform was the AT2G29630.28 type II RNA; the highest expressed minor isoforms seen during the light period are a type I isoform and another type II isoform (Additional File 2: Fig. S11C). Although the total expression profiles using AtRTD3 and AtRTD2 are very similar, the underlying transcript profiles were quite different and reflect incorrectly assembled transcripts and the absence of type II transcripts in AtRTD2 (Additional File 2: Fig. S11D). Thus, the more comprehensive transcript set in AtRTD3 along with the ability of Iso-seq to identify TES, successfully distinguished the different THIC RNA classes and showed that a type II isoform is the most abundant class [61]. The impact of increased diversity and transcript profiling resolution were also illustrated by the identification of a novel cold-induced isoform with shorter TSS and TES in AT3G17510 (CBL-INTERACTING PROTEIN KINASE 1 - CIPK1) and a novel isoform (AT4G25080.13) encoding an N-terminally truncated protein of AT4G25080 (MAGNESIUM-PROTOPORPHYRIN IX METHYLTRANSFERASE - CHLM) in AtRTD3 (Additional File 2: Fig. S12 and 13, respectively).

### Cold- and light-induced differential TSS and poly(A) site usage

Differential TSS and TES usage was observed among the expressed isoforms of AT3G17510 (CBL-INTERACTING PROTEIN KINASE 1) (Additional File 2: Fig. S12). To examine differential TSS and TES usage more widely, we applied the Time-series Isoform Switch (TSIS) program [63] to the RNA-seq cold time-course analysed with AtRTD3 as reference to identify genes with isoforms whose relative abundance switched significantly between different time-points. These isoform switches could be due to differential TSS/TES usage as well as variation in AS events. To identify genes with significant isoform switches in genes with alternative TSS and TES, we extracted lists of genes which contained alternative TSS and TES which were more than 100 bp apart (2251 and 1753 genes, respectively). To demonstrate that at least some of these genes had differential TSS usage, we compared the 2251 TSS genes to 220 genes which were previously shown to have blue light-induced differential TSS usage [54]; 82 genes were common to both groups. The TSS and TES genes lists were then used to filter the TSIS output and manual inspection of transcript isoforms distinguished switches due to alternative TSS usage (Fig. 6A-D) and TES usage (Fig. 6E, F). For example, the AT1G11280.11 isoform had a TSS 123 bp upstream of the .6 isoform and their poly(A) sites differed by only 3 nt. The .11 transcript (3,473 nt including introns) has an intron in the extended region and codes for a protein of 830 amino acids with 10 additional amino acids at the N-terminal end compared to the .6 isoform (Fig. 6A). At 20 ⁰C, the .6 isoform peaked 3 h after dusk (T2) and then declined in expression; cold rapidly induced expression of this transcript in the dark while expression of the .11 transcript does not change significantly in response to light-dark or cold (Fig. 6A). AT3G13110 is a single exon gene. The .1 and .2 isoforms have the same poly(A) sites but the TSS of .2 is 272 bp upstream of .1. The .2 transcript codes for a protein with a 55 amino acid N-terminal extension. At 20 ⁰C there was little expression of the .1 transcript but cold caused a rapid, transient increase in day 1 at 4 ⁰C peaking at dawn (T13) while the .2 transcript showed a modest increase at low temperature. Thus, at 20 ⁰C the .2 promoter drives expression and cold induces a rapid switch to the .1 promoter (Fig. 6B). The .11 isoform of AT1G55960 has a TSS 104 bp upstream of the .7 isoform and slightly different poly(A) sites (differing by 12 nt); the isoforms code for identical proteins (Fig. 6C). At 20 ⁰C, both isoforms were expressed in the light peaking 3 h after dawn (T5). However, expression levels of .11 were lower than .7 in the dark but showed a large increase in expression in the light at both 20 ⁰C and 4 ⁰C (day 1) which was lost by day 4 at 4 ⁰C (Fig. 6C). Thus, AT1G55960 has a light- and cold-regulated promoter switch. The TSS of the .12 isoform of AT5G53420 is 717 bp upstream of that of the .7 isoform. The poly(A) site of .12 is also longer by 47 nt and codes for a 265 amino acid protein including a 79 amino acid N-terminal extension (Fig. 6D). At 20 ⁰C, the shorter .7 transcript was expressed rhythmically during the day and declined in the cold with a rapid switch to higher expression of the longer .12 isoform mainly in the dark with different phasing of expression (Fig. 6D). This suggests that the promoter driving expression of the .7 transcript is light responsive and negatively regulated by low temperature while that of the .12 isoform is cold-responsive.

Differential TES usage was shown for the .26 and .27 isoforms of AT4G14400 which have identical TSS and code for the same protein but have different poly(A) sites, 194 nt apart. At 20 ⁰C, expression of .27 was significantly higher than .26 peaking at dusk (T1) while .26 peaked 3 h later in the dark (T2). Expression of the isoforms increased during the day but in day 1 at 4 ⁰C, the .26 isoform increased to a similar level to the .27 isoform (Fig. 6E). The differential phasing of expression of the isoforms was more pronounced at 4 ⁰C (Fig. 6E). The isoforms only differ by their poly(A) sites suggesting that phasing of expression and the cold response of .26 are mediated by alternative polyadenylation. Finally, the .24 and .12 isoforms of AT3G56860 have identical TSS and CDS but very different poly(A) sites with that of the .24 isoform being 1,218 nt downstream (Fig. 6F). Both isoforms were expressed at 20 ⁰C in an almost complementary way but at 4 ⁰C there was a rapid increase in expression of the shorter .12 isoform and decline of the .24 isoform. Thus, the very different cold responses of the two isoforms may be controlled by alternative polyadenylation. The TSIS method only identified a subset of potential differential TSS and TES usage because it was limited to genes which had TSS or TES sites that were > 100 bp apart and where different isoform abundances switched significantly. Nevertheless, defining TSS and TES by Iso-seq allows detailed investigation of developmental stage- and condition-specific changes in TSS and TES usage.

## Discussion

The accuracy of differential gene expression and differential alternative splicing analyses of RNA-seq data depends on the quality and comprehensiveness of the reference transcriptome. Here, we present a new Arabidopsis RTD (AtRTD3) which has extensive support from single molecule sequencing (PacBio Iso-seq). Data was generated from a wide range of organs/tissues, abiotic and biotic treatments, and RNA-processing mutants to increase the number and diversity of transcripts. Novel methods were developed to identify high confidence SJs and TES/TSSs to overcome 1) the sequencing errors particularly around splice junctions which generate thousands of false transcript structures/annotations and 2) the impact of degradation and truncated transcripts/reads on accurate end determination. In AtRTD3, 78.7% of transcripts (from 63.6% of genes) are high quality Iso-seq-derived transcripts with accurately defined SJs and start and end sites. For those genes with little or no Iso-seq coverage, transcript isoforms were taken from AtRTD2 (14.8%) and Araport11 (6.5%). AtRTD3 contains 169,503 unique transcripts from 40,932 genes reflecting novel genes (mostly lncRNA genes), novel AS transcripts and defined TSS/TES compared to the short read-derived AtRTD2 [39]. AtRTD3 represents a high quality, diverse and comprehensive transcriptome which improves gene and transcript quantification for differential expression and AS analysis and now allows alternative TSS and TES usage to be addressed.

In the production of AtRTD3 we applied a hybrid analysis pipeline using PacBio Isoseq3 and TAMA and developed new methods of single molecule sequencing analysis which are generally applicable and will improve downstream analysis and the quality of transcript and transcriptome annotations. We showed previously that redundant or missing transcripts, transcript fragments, and variation in the 5’ and 3’ ends of transcripts of the same gene seriously impacted the accuracy of transcript and gene expression quantification with Salmon and Kallisto which require prior knowledge of transcripts [39]. Initial analysis of the Iso-seq data identified issues with false splice junctions, degraded or fragmentary reads/transcripts and that error correction methods using short read data often trim or split whole transcripts sequences in fragments or generated new errors (over-correction). In addition, the Isoseq3 analysis pipeline from PacBio used polishing steps which removed splice site variation with small differences such as alternative splicing of a few nucleotides (e.g. NAGNAG sites). These observations provided the motivation to improve methods of analysis of PacBio Iso-seq data. Firstly, we used the Isoseq3 pipeline up to the generation of FLNCs and then switched to TAMA which gave greater control over transcript processing and was the basis of developing the SJ centric approach. Secondly, we clearly demonstrated that mismatches in the vicinity of SJs generated transcripts with false splice junctions. We defined criteria to identify high confidence splice junctions and remove poorly supported SJs. The number of rejected SJs and the high overlap with the accurate short read-determined SJs illustrated the value of the splice junction centric approach. Thirdly, even with 5’-cap capture, there is extensive variation in transcript start and end sites, much of which reflects degradation of RNA. Distinguishing high confidence TSS and TES from such degradation products required different methods that take into account the effects of different gene expression levels and the stochastic nature of transcription start sites. The high confidence TSS and TES defined in AtRTD3 were supported by the frequency, position and distribution of conserved promoter, polyadenylation and translation start motifs and by good agreement with experimentally defined TSS and poly(A) sites [41,43,44]. Such experimental determinations are often limited in the number of genes for which data is generated and the number of transcripts where both the 5’ and 3’ ends are defined. The new pipeline addresses the major issues of accuracy of splice site and TSS/TSS determination in Iso-seq analysis. The methods have three main advantages: 1) the generation of high confidence SJs removed the need for error correction using short reads and therefore avoided splitting or trimming of the original sequences as well as over- correction, 2) both TSS and TES are generated for a very high proportion of transcripts, and 3) they are determined directly from the single molecule data without the need for parallel experimental approaches. To date, Iso-seq has been applied to a wide range of plant species (see Background); the novel methods here will improve analysis of transcripts in future studies and allow re-analysis of existing data. In addition, AtRTD3 can evolve further with the addition of new or existing Iso-seq datasets analysed using the methods described here.

The Iso-seq derived transcripts in AtRTD3 (ca. 80% of transcripts) were full-length with accurate SJs and TES/TSS and correct combinations of TES/TSS and AS events but only covered ca. two-thirds of genes in Arabidopsis. This represents good coverage for Iso-seq in comparison to other studies. For example, a recent study of Iso-seq of nine tissues in rice covered only ca. one-third of rice genes [36]. Coverage of the other genes and transcripts in AtRTD3 came from Araport11 and, primarily, from AtRTD2 due to its far greater transcript diversity [39]. The transcripts from AtRTD2 and Araport11 are of high quality in terms of splice sites but their 5’ and 3’ ends are likely to be inaccurate and are often artificially extended [39]. The quality of SJs in the AtRTD2 transcripts is evidenced by 57.8K of the 82K AtRTD2 transcripts being redundant to Iso-seq transcripts in having identical SJs such that the Iso-seq transcripts were preferentially selected. Thus, AtRTD3 has full coverage of the genes in Arabidopsis with two-thirds of genes made up predominantly of Iso-seq transcripts and one-third of high quality RNA-seq assembled transcripts. AtRTD3 is unique in that all of its transcript annotations have undergone extensive quality controls. As higher accuracy and throughput of single molecule sequencing technologies improve, the new analysis pipeline exploited here will enable the rapid determination of SJs, TSS and TES for fully comprehensive transcriptomes.

AtRTD3 contains greatly increased numbers of unique transcripts and particularly transcripts coding for protein variants and unproductive transcripts from protein-coding genes compared to AtRTD2. Although transcript numbers more than doubled in AtRTD3, 60.4% of multi-exonic protein-coding genes had AS agreeing with previous estimates [39, 58]. The increased number of protein variant transcripts include transcripts from the same genes with alternative TSS and pA sites and the identification of novel AS events which alter coding sequences. The increased unproductive transcripts also included transcripts with the same PTC-generating AS event but with alternative TSS and TES sites and the majority contained classic NMD characteristics. Iso-seq identified novel AS events and, in particular, high numbers of intron retention events. The majority of transcripts with intron retention most likely reflect partially spliced pre-mRNAs and why such transcripts should be more prevalent in Iso-seq is unknown but may be due to lower efficiency of obtaining full short read coverage of introns in short read assembly. In plants, transcripts with intron retention have been shown to avoid NMD and to be retained in the nucleus [53, 64]. In contrast, human intron retention transcripts are generally degraded by the NMD pathway [65] but numerous examples of intron retention as a regulatory mechanism have been described [66]. For example, intron detention where partially spliced transcripts remain in the nucleus until required and are then spliced and mRNAs exported and translated represent novel gene regulation mechanisms [66]. In this regard, we have identified ca. 20K protein-coding transcript isoforms with AS only in the 5’ and/or 3’ UTR such that isoforms coded for the same protein. AS in UTRs can be involved in regulation of expression by introducing short or over-lapping uORFs to trigger NMD or affecting translation [53] or nuclear retention of mRNAs determining export of mRNAs [66]. The detailed characterisation of such transcripts here provides a basis for future investigation into the regulatory roles of AS in UTRs.

The power of exploiting comprehensive RTDs in analysing differential expression and differential alternative splicing was demonstrated in Arabidopsis using a cold time-series dataset and AtRTD2 [59, 60]. Thousands of genes with rapid cold-induced significant changes in expression and AS were identified due to the transcript level resolution of expression [59, 60]. AtRTD3 is more comprehensive and for most transcripts (ca. 80%) there is detailed structural information in terms of AS events and TSS/TES which increase the resolution of the analysis. Direct comparison of transcript quantification using AtRTD2 and AtRTD3 showed an increase in accuracy and the impact of missing transcripts and incorrectly assembled transcripts as seen previously [39]. More importantly, the defined TSS and TES clearly demonstrated variation in TSS and TES for many genes and re-analysis of the cold time-series data with AtRTD3 identified differential TSS and TES usage due to low temperature and light/dark conditions. It will now be possible to examine transcriptional and post-transcriptional regulation of gene expression involving differential TSS and TES usage demonstrated here and the impact of AS in UTRs [53, 67] during development and in response to abiotic and biotic stresses. Differential TSS and TES usage illustrates novel regulatory mechanisms. For example, Kurihara et al. [54], identified differential TSS usage in response to blue light and proposed a mechanism whereby blue light induces use of a TSS downstream of an uORF to produce a transcript that avoids NMD and allows expression. As mentioned above, over 20K transcripts in AtRTD3 have AS only in the UTRs and interplay between TSS/TES usage and AS in the UTRs may have important regulatory roles affecting stability of transcripts, whether they are retained in the nucleus or exported, and avoid NMD or are degraded to fine tune gene expression.

## Materials and Methods

### Plant material

Plant samples for RNA extraction and Iso-seq sequencing were all from Arabidopsis Col-0 and are summarised in Additional File 1: Table S1 and described below.

#### Different organ samples

flower, silique and root materials. Col-0 was used for all samples. Roots: roots were harvested from 5-week-old plants grown in liquid culture (12 h light/12 h dark) and harvested at dawn and dusk and pooled. Siliques and inflorescence/flowers: plants were grown in soil in 16 h light/8 h dark conditions at 23 ⁰C; siliques of different sizes (stages) up to early browning and inflorescences containing flowers from buds to mature flowers were harvested from 6-week-old plants and each pooled. For etiolated seedling samples, seedlings were grown for 3, 4, 5 and 6 d in darkness on petri dishes (½ Murashige and Skoog medium) without sugar and samples were pooled.

#### Plants exposed to different abiotic stresses/cues

Cold, heat, flood and time-of-day. Cold: 5- week-old rosettes grown in 12 h light/12 h dark and 20⁰C were exposed to 4⁰C at dusk for different lengths of time (12 h and 66 h) and samples were pooled; Heat – 5-week-old rosettes and 12-day-old seedlings grown in 16 h light/8 h dark at 23⁰C and 20⁰C, respectively, were exposed to high temperatures (27⁰C and 37 ⁰C, respectively) for different lengths of time (1 week and 12 h, respectively), harvested (4 h after dawn) and pooled; Flooded: 5-week-old rosettes grown on soil with 16 h light/8 h dark at 23⁰C were either flooded or completely submerged under water for two different time exposures (24 h and 6 d) and pooled; time-of-day – 5-week-old rosettes were grown under 12 h light/12 h dark at 20⁰C and were harvested at dawn and 6 h after dawn.

#### UV-C treatment

Col-0 seedlings were grown on ½ Murashige and Skoog agar plates at 22 °C under 12 h light/12 h dark conditions until the first pair of true leaves was expanded (9 d after germination). The ultraviolet treatment was performed using a Stratalinker (Stratagene) at 254 nm with 1 kJ/m². Subsequently, seedlings were incubated in either light or dark. Whole seedlings were collected after 1 and 4 h of incubation and frozen in liquid nitrogen. Equal amounts of RNA from UV-C treated samples were pooled.

### Plants infected with different pathogens

#### Botrytis cinerea, Hyaloperonospora arabidopsidis, and Pseudomonas syringae

For B. cinerea infection, detached 5-week-old Arabidopsis (Col-0) leaves (grown at 22°C, 12h light/12 h dark, 60% humidity) were placed on agar, and inoculated with 5 x 7 uL droplets of 100,000 spores per mL in 50% grape juice. Infected trays were sealed and kept at 22°C, 12h light/12 h dark, 80% humidity. Samples (two infected leaves) were collected by flash freezing in liquid nitrogen at 24 h, 30 h and 36 h post-inoculation . For H. arabidopsidis infection, 14-day-old Col-0 seedlings (grown at 22°C, 12h light/12 h dark) were sprayed with 30,000 spores per mL in water of Hpa isolate Noks1, 15 mL per P 40 tray (0.375 mL per module), sealed and at grown at 18°C, 12h light/12 h dark. Infected seedlings were harvested s at 4, 5 and 7 days post-inoculation and flash frozen in liquid nitrogen. RNA was extracted and RNA samples pooled within each pathogen (final pool included 2 samples per time point). For P. syringae infection, 3-week-old plants were infected with P. syringae pv tomato DC3000 by infiltrating three leaves of five plants with 2x10^5^ cfu/ml at ZT2 (12 h light/12 h dark). Infiltrated leaves were harvested 8 h and 24 h post-infiltration. RNA was extracted from both time-points and pooled.

### Material from RNA processing/degradation mutants (NMD and exosome) and nuclei

Mutants were an NMD double mutant combining the heterozygote of lba1 (upf1) and knockout upf3-1 and exosome mutants: xrn3-3, xrn4-6 and xrn2-1. Seedlings were grown on petri dishes and those of the exosome mutants pooled together. Nuclei were prepared from leaves of 5-week-old plants.

### RNA extraction and library construction

For the majority of samples, RNA was isolated with the RNeasy plant mini kit (QIAGEN – including on-column DNase I treatment) according to the manufacturer’s instructions. RNA was extracted from etiolated seedlings, the NMD double mutant and nuclear extracts with the Universal RNA purification kit (EURx). PacBio non-size selected Iso-seq libraries were constructed using Lexogen Teloprime, Teloprimev2 or Clontech kits following manufacturer’s instructions (Additional File 1: Table S1). Each of the 27 libraries were sequenced on a single SMRT cell (1M,v3 for Teloprimev2 and Clontech) on a PacBio Sequel machine using a 10hr (v3) movie.

### Analysis of PacBio Iso-seq reads

The workflow of the analysis is shown in Fig. 1. The PacBio sequencing data was analysed using the PacBio Isoseq 3 pipeline to generate and map full-length non-chimaeric (FLNC) reads. Further analysis was performed using TAMA [7] (https://github.com/GenomeRIK/tama) to collapse and merge reads/transcripts and apply novel methods to define splice junctions (SJs) and transcript start and end sites (see below).

### Processing of raw PacBio IsoSeq reads to FLNCs

The raw PacBio sequencing data (.subreads.bam) from each library was processed individually using the following procedures: 1) CCS calling was carried out using ccs 4.0.0 using the following parameters: --min-rq 0.9 -j 28. 2) Primer removal and demultiplexing was carried out using lima (version v1.10.0) with the parameters: --isoseq --peek-guess; 3) isoseq3 (v3.2.2) refine was used to trim poly(A) tails and for rapid concatemer identification and removal to produce the FLNC transcripts (Fig. 1A). For the Clon tech libraries, --require-polya is used while for Teloprime 5’ captured reads, lima is run with this parameter turned off. We have deliberately avoided the clustering steps in the Isoseq3 pipeline in order that small variances around the splice junctions, such as NAGNAG splice junctions can be preserved. The FLNCs were then converted to FASTA format using samtools and mapped to the TAIR10 genome reference using minimap2 (version 2.17-r941) using the following parameters -ax splice:hq -uf -G 6000. The mapping files (bam files) were then sorted and the non-mapped reads were filtered out.

### Splice junction centric approach for accurate splice junctions

From this point, we adopted the TAMA analysis pipeline for the next steps of transcript isoform analysis (Fig. 1B). To overcome the generation of false splice junctions due to mis-mapping of FLNCs to the genome, we developed a splice junction centric approach to provide highly accurate alignment around splice junctions. A n improvement of TAMA was developed that allowed us to examine the mapping mismatches (replacement and indels) between the FLNCs and genome reference. Using this new parameter (-sjt and -lde), we were able to extract the mapping details of any defined regions around the SJs. For each library, we ran the TAMA collapse using the following parameters “-d merge_dup -x no_cap -m 0 -a 0 -z 0 -sj sj_priority -lde 30 -sjt 30” so that 1) small variations of up to 1nt at SJs, as well as transcription start and end sites, are preserved in the FLNC reads; 2) mapping details of 30 nt around each SJ were extracted. Then TAMA merge (merged -m 0 -a 0 -z 0 -d merge_dup) was used to merge all the transcripts from the libraries and all the redundant FLNCs were removed, while the small variations up to 1nt at SJs, as well as transcription start and end sites, were preserved in the merged FLNC reads. To accurately determine splice junctions, we examined the high-resolution alignment information around the SJs and found that high confidence SJs are always supported by at least one alignment with a perfect match between the FLNCs and the genome reference around the SJ. SJs were also compared to those of AtRTD2 and their sequences assessed using Position Weight Matrix - PWM) [68]. To derive a list of high confidence SJs (Fig. 1C) (and thereby identify falsely aligned SJs), our SJ centric approach employed the following criteria: 1) the presence of canonical splice junction motifs; and 2) no mapping mismatches including substitution, deletions and insertions, with 10nt around the SJs with support of at least one read.

### Determination of transcription start and end sites

For high abundance genes, we assume that Iso-seq reads with authentic TSS/TES sites would be sequenced more often than those representing degradation products where end sites will occur randomly. We can use the binomial distribution to estimate the probability of having m Iso-seq reads underpinning one specific start by random.

For m Iso-seq read starts at n genomic locations, with the assumption that the starts of the degraded Iso-seq reads are random, we assume the probability of each read to have a start at particular genomic location (p) is equal among all the read start locations, thus 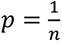. The probabilities of having k reads at one genomic location at random can be calculated as a bionomial probability 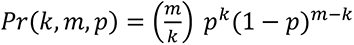. A smaller probability would indicate that the start/end genomic location is unlikely to have such a number of reads (low and high) at random. We are interested in identifying the non-random start locations that have higher numbers, so we have applied the following criteria: 1) k should be higher than the average reads for all genomic locations for that gene 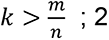) the probability of having k of reads at one genomic location should be small with *Pr*(*k*, *m*, *p*) < 0.05.

We define the 5’ location of the long read as RSGLs and 3’ location as REGLs. The non-random RSGLs and REGLs with higher than expected numbers of reads are defined as significant RSGLs and REGLs, which are likely to be TSS/TES sites. Additionally, we removed REGLs which could be a result of off-priming identified by the REGLs being followed by poly(A) sequences in the genome.

For low abundance genes where we could not detect significant RSGLs and REGLs, we applied a different set of criteria. Reads were compared and a significant start or end site required at least two long reads supporting that site within a sliding window of 11nt (5nt on each side).

To account for the stochastic nature of the TSS/TES, a 100nt window around significant RSGLs and RSGLs were defined as high confidence TSS/TES regions. All the merged FLNCs from all of the libraries were then filtered based on the high confidence SJs and high confidence TSS/TES regions (Fig. 1C). Transcripts containing SJs, TSS and TES which did not match the high confidence set were removed. To generate high level transcripts, transcripts with small variances in 5’ and 3’ UTR lengths were removed by further collapsing transcripts by running the TAMA merge on the filtered FLNCs using “-m 0 -a 50 -z 50 -d merge_dup” that allows transcripts with variations within 50 nt at UTR regions to be merged. Thus, to achieve accurate transcript isoforms from the PacBio data and generate AtIso (Fig. 1C), we have adopted a strategy that seeks evidence to support all SJs, TSSs and TESs. Finally, to increase the gene coverage using existing annotations and make the maximum use of the Iso-seq long reads, we retained genes that overlapped with Araport annotation on the same strand (>50%). These were combined with the genes with TSS/TES support to generate the final set of genes and transcript in AtIso.

### TSS and TES motif enrichment analysis

To search for known TSS/TES related motifs around significant RSGL and REGLs as well as the identified loci of interest in other datasets (potential TSS and TES sites), the following approach was taken. A number of motifs associated with TSS and TES sites were identified (Additional File 1: Table S6). For each identified TSS and TES, the sequence within ±500 nucleotides on each side was extracted from the genome. A regular expression search was carried out in the extracted sequences searching for the known enriched motifs related to TSS and TES. All matching motifs and their positions relative to the site of interest were extracted. From this the number of instances of the motif were calculated for every position ±500 nucleotides relative to the TSS/TES. As a control, the same number of random sites were taken, and the above analysis was carried out.

### Construction of AtRTD3

AtIso represents the most accurate and extensive representation of Arabidopsis transcripts to date. To overcome the low coverage of genetic regions and the lack of transcript diversity in genes with low expression, we integrated the transcripts from short read assemblies AtRTD2 and Araport into AtIso to generate the comprehensive transcriptome, AtRTD3. In AtRTD3, we kept all the transcripts from AtIso and only introduced transcripts from AtRTD2 and Araport that 1) contained novel SJs (AtRTD2 and Araport) or 2) covered genomic loci in Araport not covered by Iso-seq. The novel SJs were identified in a pairwise fashion in sequential order by, firstly, comparing AtIso and AtRTD2, extracting the transcripts in AtRTD2 with novel SJs that were not in AtIso, and, secondly, repeating the process with transcripts in Araport containing unique SJs (not in AtIso and AtRTD2). The transcripts from Araport covering novel loci that did not overlap with AtIso are also extracted. Finally, all the extracted transcripts mentioned above were merged together with AtIso using TAMA merge (-m 0 -a 50 -z 50 -d merge_dup).

During merge, we give Iso-seq assembled transcripts the highest priority by setting the “cap_flag” as “capped” and “merge_priority” as “1,1,1”, indicating 5’ TSS, splice junctions as well as 3’ TES of Iso-seq assembly all take highest priority during merging. For short-read assemblies we label “merge_priority” as “uncapped” and “merge_priority” as “2,1,2” . This means that only the SJs were given top priority as they have been validated by short reads. 5’ TSS and 3’ TES from the short-read assembly would be lower priority and contribute less to the determination of the TSS and TES when merging with Iso-seq transcripts.

### Annotation of AtRTD3

To annotate AtRTD3, we examined the overlaps of AtRTD3 transcripts with Araport gene annotations using bedtools (intersect -wao). Transcripts were assigned to the Araport genes if they overlap on the same strand (where the overlap covers >30% of both transcripts). Transcripts that overlap two Araport genes on the same strand would be assigned a gene ID with two concatenated gene names (e.g. AT1G18020-AT1G18030). This allows the identification of biological chimeric transcripts that run-through two or more genes. The origin of these transcripts (AtIso, AtRTD2 or Araport11) are also added in the bed annotation to allow users to distinguish high confidence transcripts from long read assemblies from less confident transcripts from short read assemblies.

### Availability of data and materials

All the sequencing data in this study have been deposited in the Short Read Archive (SRA) (https://www.ncbi.nlm.nih.gov/sra) with BioProject ID: PRJNA755474. The atRTD3 annotations are available in fasta, bed and gtf format at https://ics.hutton.ac.uk/atRTD/RTD3/ (username atrtd; password atrtd3-06092021). The script that used to carry out the analysis in this manuscript can be found at https://github.com/ZhangTranscriptomislab/atRTD3

## Contributions

R.Z., J.B., A.S.N.R., A.B., M.K. designed and managed the project. C.P.G.C, S.R., S.H., A.M. and M.T. collected the samples and extracted RNA. R.Z., R.K. and M.C. developed the methods for Iso-seq analysis, Y.G. and L.G. performed initial analysis of the data before development of new methods. J.C.E. developed TranSuite, Y.M. generated the PWM data, C.P.G.C. carried out correlation analyses and W.G. analysed the cold time series data with AtRTD3 and TSIS. R.Z. and J.B. performed data analyses, assessed outputs and interrogation of results. J.B. and R.Z. organized the data and wrote the manuscript. L.M. created hosting website for AtRTD3. The groups K. D./S. H., M. K./A. B./S. R., D. S./T. K., M. S./A. M./M. T., A. W./T. W.-K. and J. B./C. C. provided grew plant samples and provided RNA; M. C., K. D., A. b. H., E. H., M. J., A. J., T. K., S. L., Q. Q. L., T. M., M. S., D. S., R. S., Z. S., S.-L. T., T.W.K., A. W., R. W., L. X., X.-N. Z., A.S.N.R., A.B., M. K. and J. B. provided samples and were responsible for funding. All authors read and approved the final manuscript. We also thank Tai Montgomery, Colorado State University for help and support.

## Corresponding author

Correspondence to Runxuan Zhang.

## Ethics declarations

Ethics approval and consent to participate Not applicable.

## Supplementary Information

Additional file 1: Table S1. Plant material for RNA samples for Iso-s eq. Table S2. Read statistics for Iso-seq libraries. Table S3A and 3B. Number and percentage of splice junctions with a sequencing error in positions L1 to L30 for A) upstream (left) and B) downstream (right) of splice junctions. Table S4. Position Weight Matrix scores for introns. Table S5A and 5B. Filtering of SJs on the basis of mismatches in each nt position. Table S6. Sequence motifs for validation of TSS and TES sites Table S7. AtRTD3 - Transcript characteristics and translations from TransFeat. Table S8A. TranSuite output of comparison of AtRTD2 and AtRTD3 for mono-exonic/multi-exonic genes with single or multiple transcript isoforms. Table S8B. Comparison of TranSuite output of AtRTD2 and AtRTD3 gene and transcript characterisation. Table S9A. AtRTD3 - novel genes. Table S9B. AtRTD3 - Chimeric genes and transcripts. Table S10. Frequency of AS event type among AtRTD3, AtIso and Araport11.

Additional file 2: Fig. S1. Iso-seq transcripts contain false SJs prior to accurate SJ determination and filtering. Fig. S2. Determination of TSS and TES sites for genes with high and low abundance reads. Fig. S3. Comparison of AtIso TSSs and TESs with previously published transcript start and end sites. Fig. S4. Generation of high level transcripts. Fig. S5. Increased number of transcript isoforms in AtRTD3. Fig. S6. Genes with characterised different TSS. Fig. S7. Genes with characterised alternative TES. Fig. S8. Confirmation of AS variant isoforms in the lncRNA, FLORE. Fig. S9. Novel genes in AtRTD3 – lncRNAs. Fig. S10. Chimeric transcripts. Fig. S11. Accurate Iso-seq transcript determination identifies THIC RNAs produced by riboswitch. Fig. S12. Novel cold-induced isoform of CIPK1 from AtRTD3.

Fig. S13. Novel transcript isoform in AtRTD3 affects expression levels of main transcripts compared to AtRTD2. Additional file 3: Figure S14. Comparison of AtIso transcripts and SJs to Araport and AtRTD2.

