## Supplementary material for "A high resolution single molecule sequencing-based Arabidopsis transcriptome using novel methods of Iso-seq analysis": Supp Figures

**Additional File 2 : Fig. S1 to S13**

**for**

A

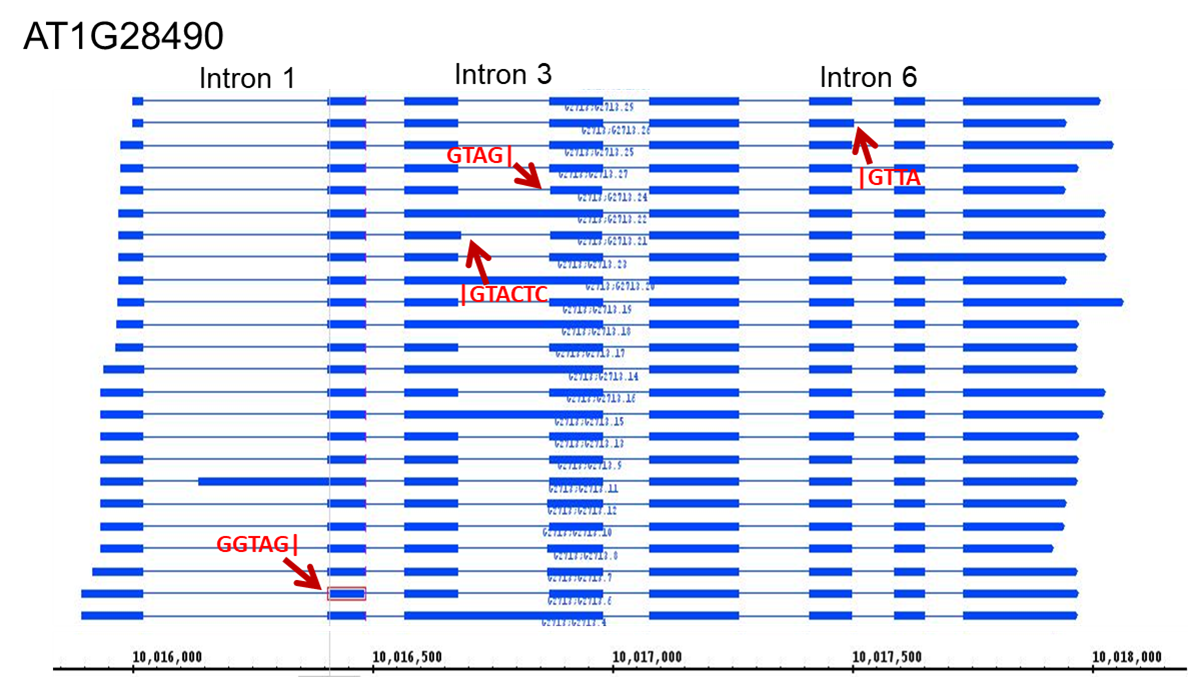

B

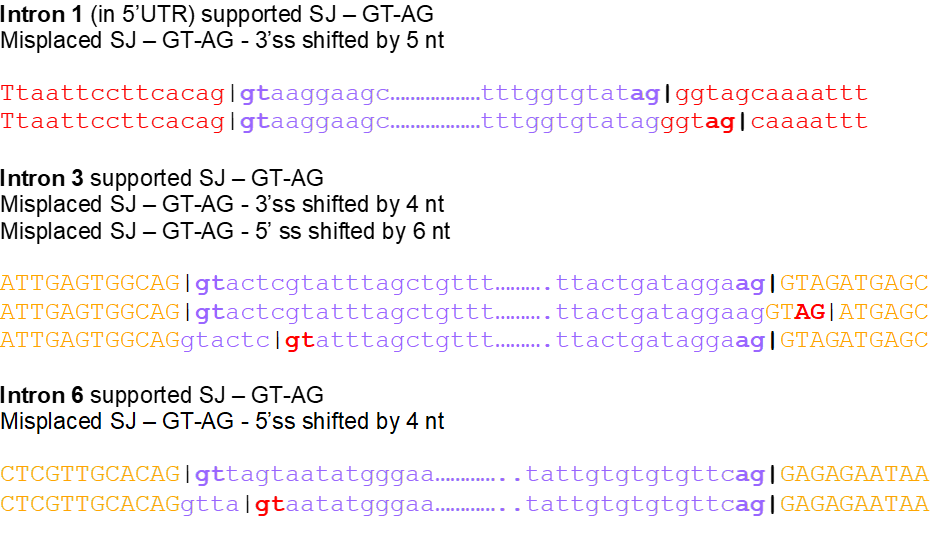

**Fig. S1. Iso-seq transcripts contain false SJs prior to accurate SJ determination and filtering. A)** Screen shot of AT1G28490 transcripts in Integrated Genome Browser. Four transcripts contain SJs which are unique to Iso-seq (not present in short read assemblies) and were not supported by an Iso-seq read with zero mismatches in the vicinity of the SJ. The mis-mapped SJs are in introns 1, 3 and 6. **B)** Alignment of authentic, supported SJs of introns 1, 3 and 6 (top line) and mis-mapped SJs showing shift in position of splice sites. 5’ UTR exon sequences – red; coding exon sequences – yellow; intron sequences (blue); authentic splice site dinucleotides – blue bold; misplaced, unsupported splice site dinucleotides – red bold.

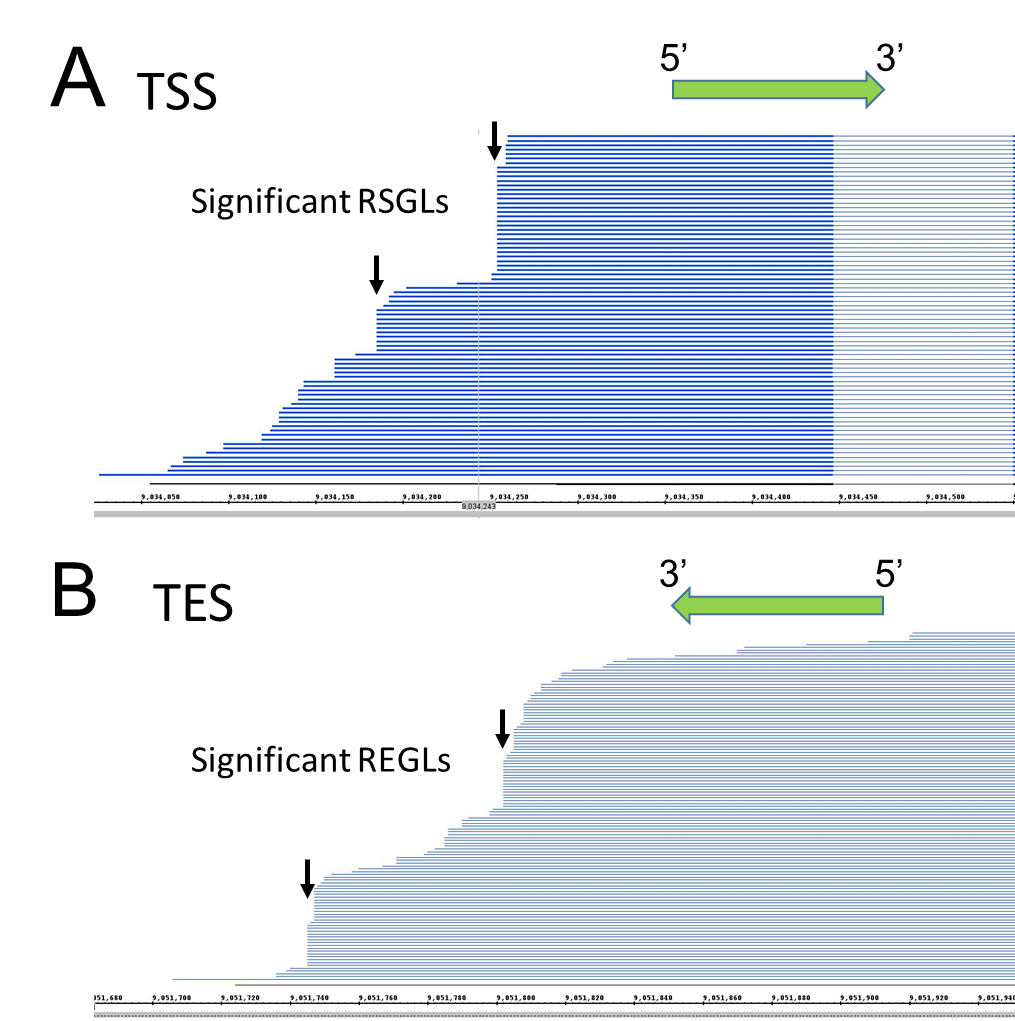

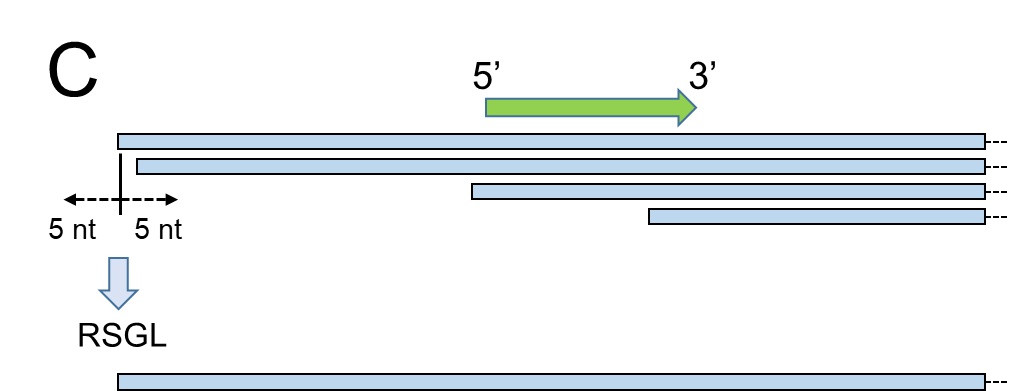

**Fig. S2. Determination of TSS and TES sites for genes with high and low abundance reads.** **A)** 5’ end of gene with multiple reads. Binomial distribution determines two significant RSGLs/TSS (arrows). **B)** 3’ end of gene with multiple reads. Binomial distribution identifies two significant REGLs/TES (arrows) on the basis of number of reads ends at specific sites. **C)** 5’-end of gene with low number of reads. Two reads with 5’-ends within an 11 nt window provide support for a RSGL.

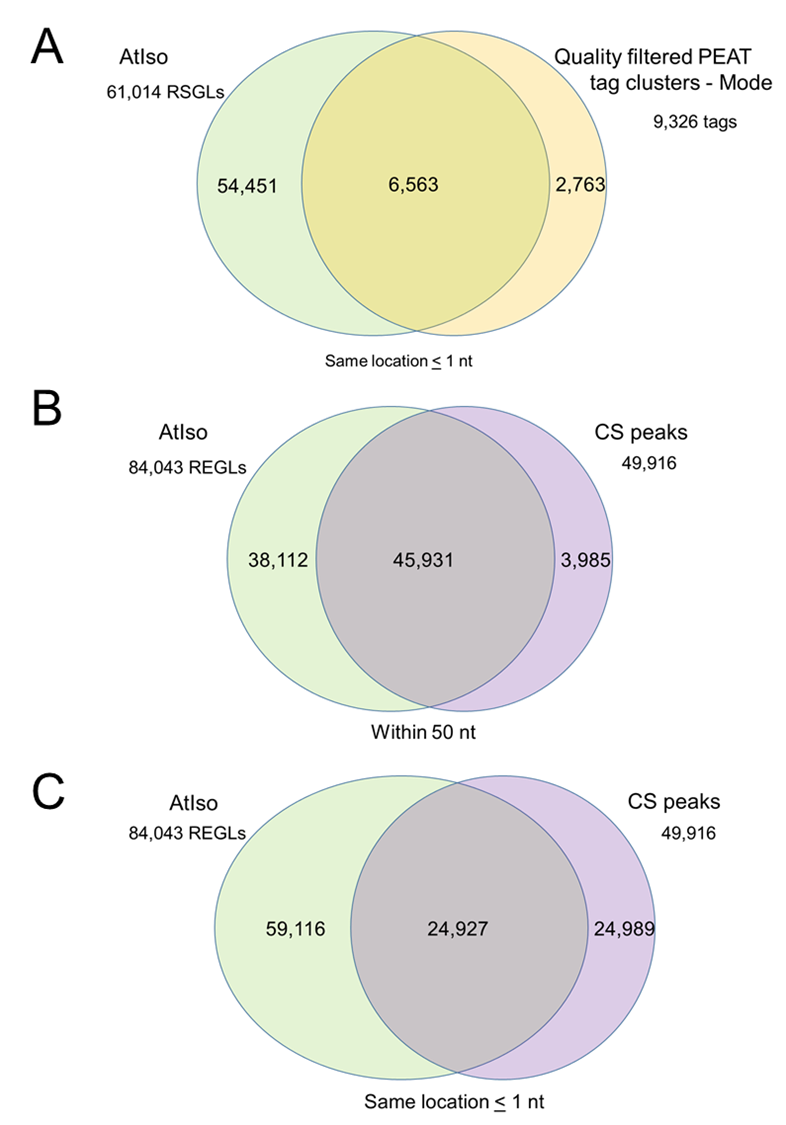

**Fig. S3. Comparison of AtIso TSSs and TESs with previously published transcript start and end sites.** **A)** AtIso RSGLs compared to the mode position of start sites from quality filtered tag clusters (within ≤ 1 nt) from Morton et al. (2014); **B, C)** AtIso REGLs compared to CS peaks from Sherstnev et al (2012) within 50 nt (**B**) and ≤ 1 nt (**C**).

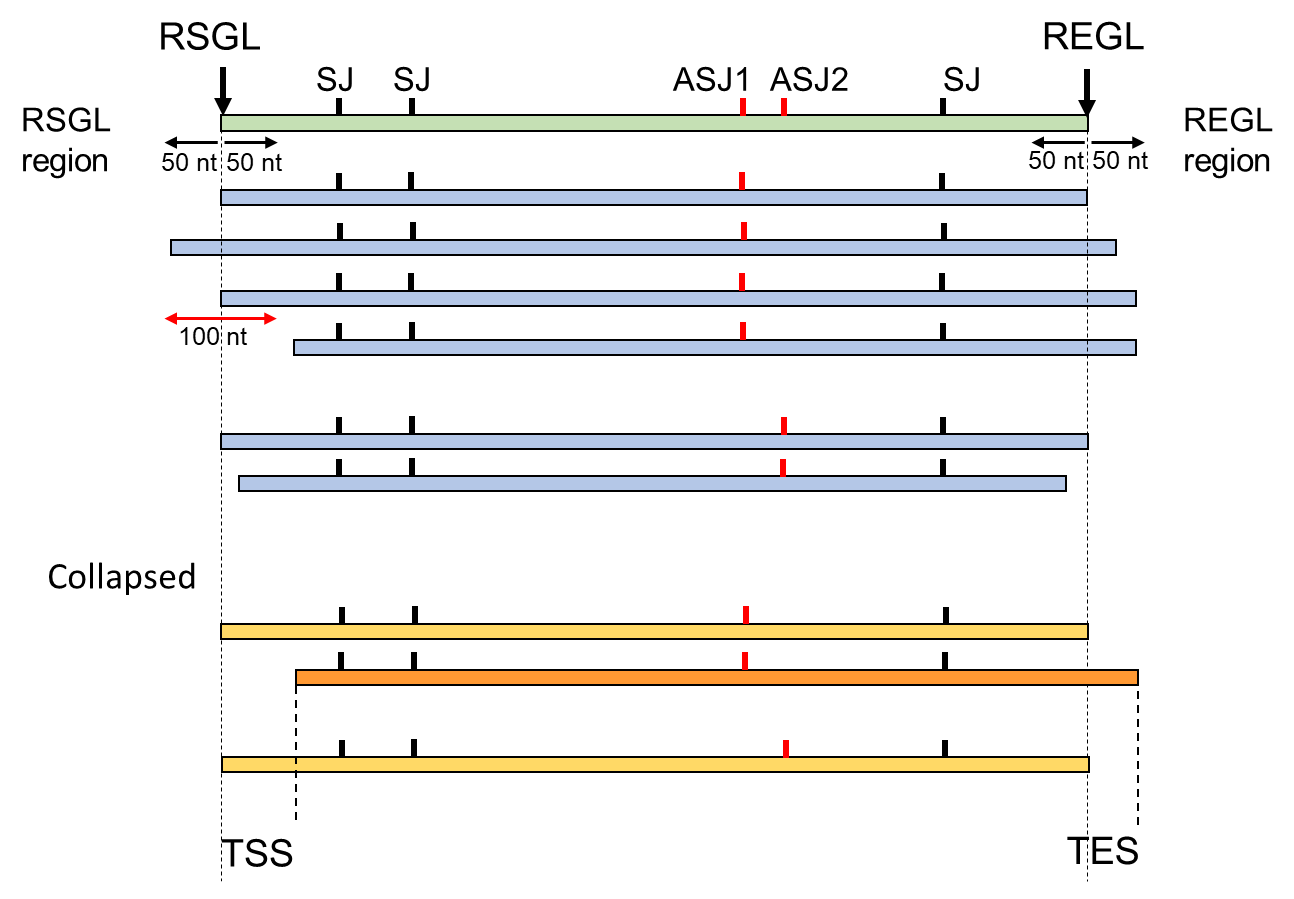

**Fig. S4. Generation of high-level transcripts.** A gene with significant RSGL/REGL positions and two alternatively spliced SJs (green). Four transcripts contain ASJ1 and two contain ASJ2 with varying start and end sites (blue). If the ends of the transcripts are within the 100 nt window of the RSGL and REGL, they are collapsed to the most common end thereby preserving the RSGLs and REGLs as TSSs and TESs (yellow) and transcripts with different TSS and TES are retained (orange). SJ – splice junction; ASJ – alternative splice junction; TSS – transcription start site; TES – transcription end site.

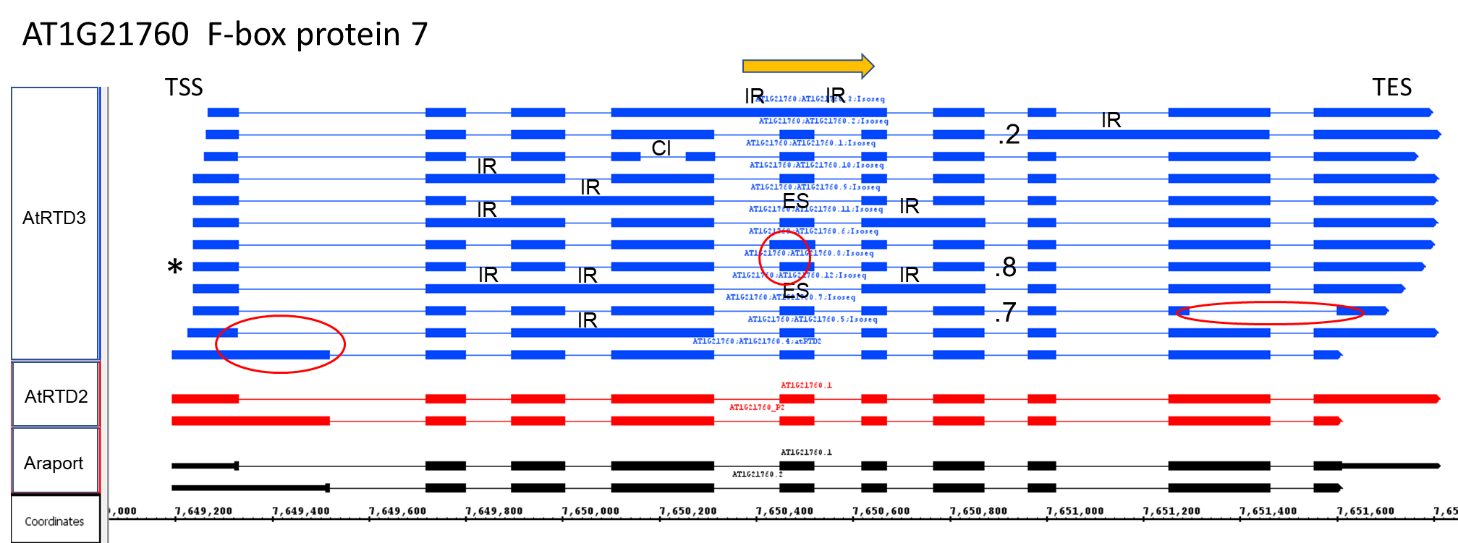

**Fig. S5. Increased number of transcript isoforms in AtRTD3.** At1G21760 has two transcripts in Araport and AtRTD2 but twelve in AtRTD3. Additional transcripts contain different AS events: intron retention (IR), exon skipping (ES), cryptic intron (CI), alternative 5’ or 3’ splice sites (circled in red) and different TSS and TES. The AT1G21760.8 transcript isoform (asterisk) codes for the full-length protein, .2 and .7 have different C-terminal ends due to AS events towards the 3’ end of the transcripts and the remaining isoforms are unproductive containing PTCs. Transcript structures visualised with Integrated Genome Browser (IGB) are from Araport (black), AtRTD2 (red) and AtRTD3 (blue); arrow shows direction of transcription.

A

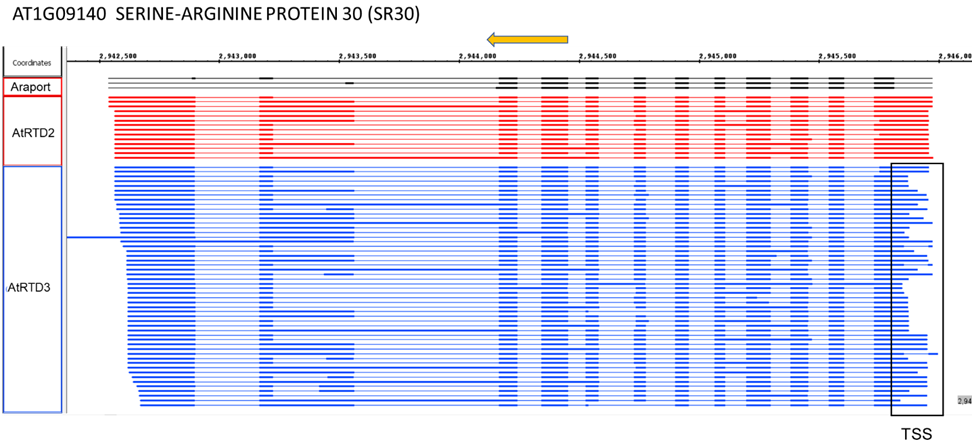

B

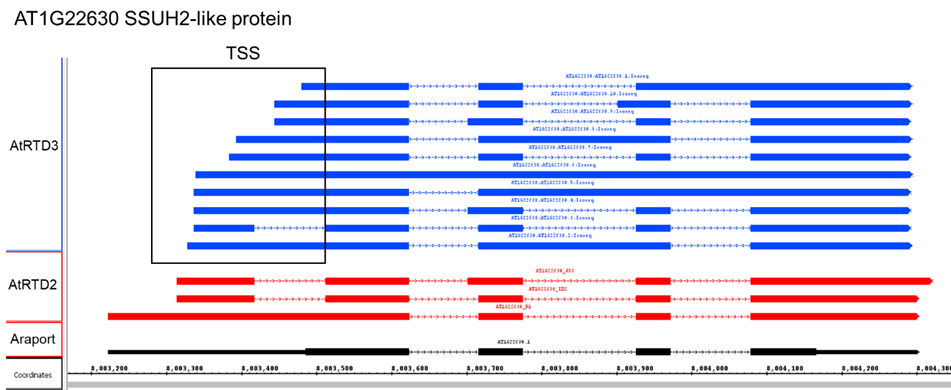

**Fig. S6**. **Genes with characterised different TSS.** **A**) AT1G09140 - AtRTD3 has 51 Iso-seq transcript isoforms reflecting combinations of different TSS, TES and AS events. **B**) AT1G22630 – 10 Iso-seq transcript isoforms with variable TSS. Both genes show differential TSS usage in response to blue light (Kurihara et al., 2018). Transcript structures visualised with IGB are from Araport (black), AtRTD2 (red) and AtRTD3 (blue); arrow shows direction of transcription.

A

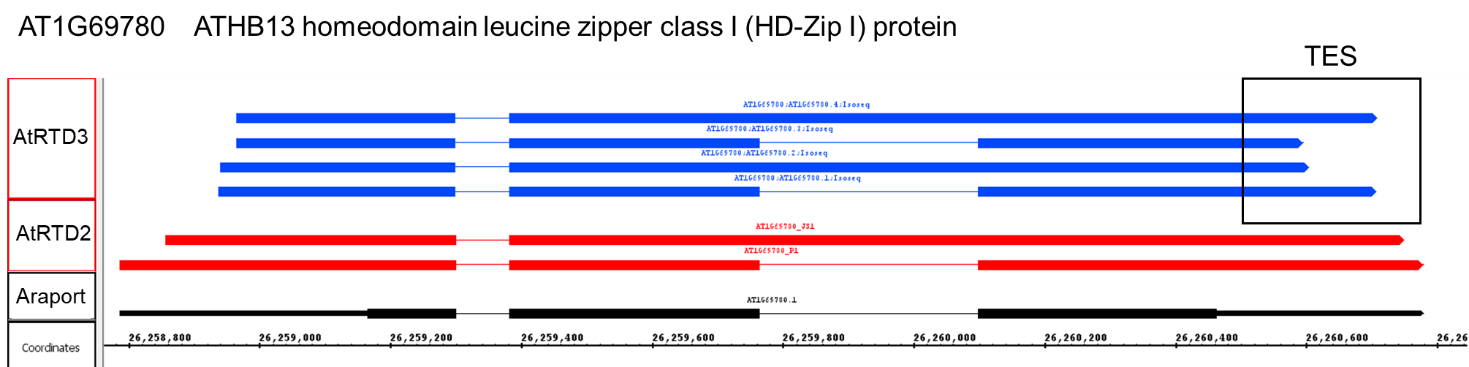

B

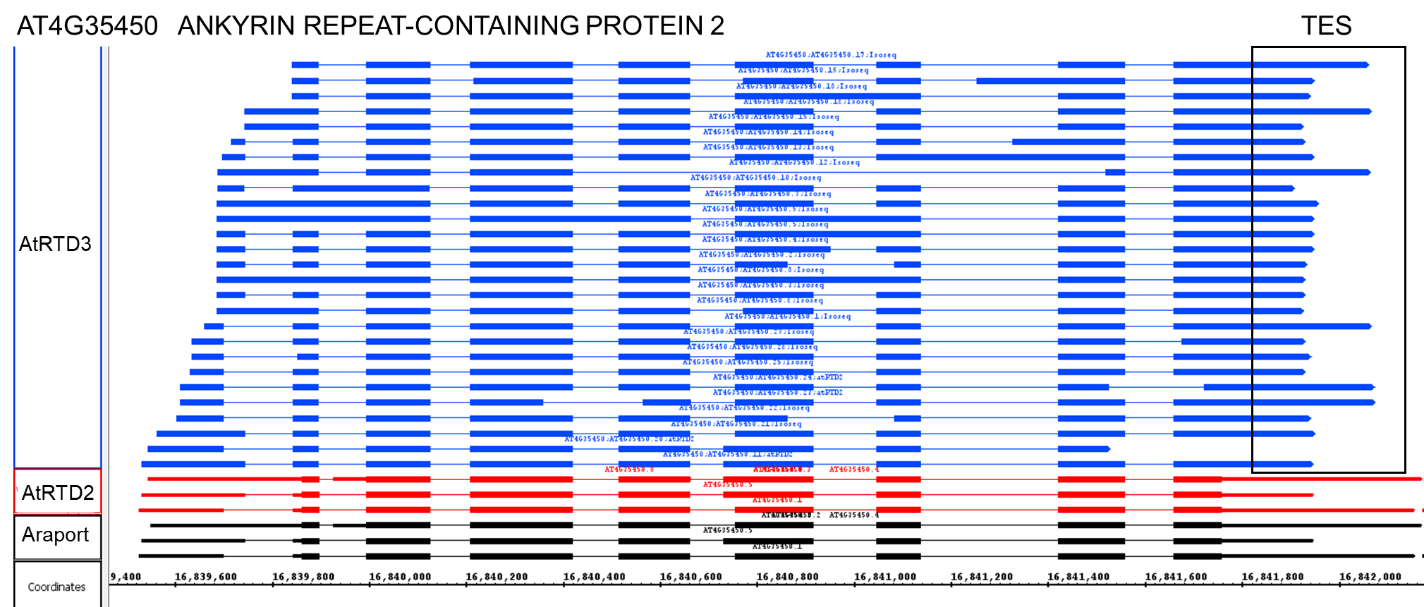

**Fig. S7.** **Genes with characterised alternative TES.** **A**) AT1G69870 - AtRTD3 has 4 Iso-seq transcript isoforms reflecting showing different TES. **B**) AT4G35450 – multiple Iso-seq transcript isoforms with variable TSS and TES. Different TES/poly A sites have been characterised previously (Yu et al., 2019). Transcript structures visualised with IGB are from Araport (black), AtRTD2 (red) and AtRTD3 (blue); direction of transcription is left to right.

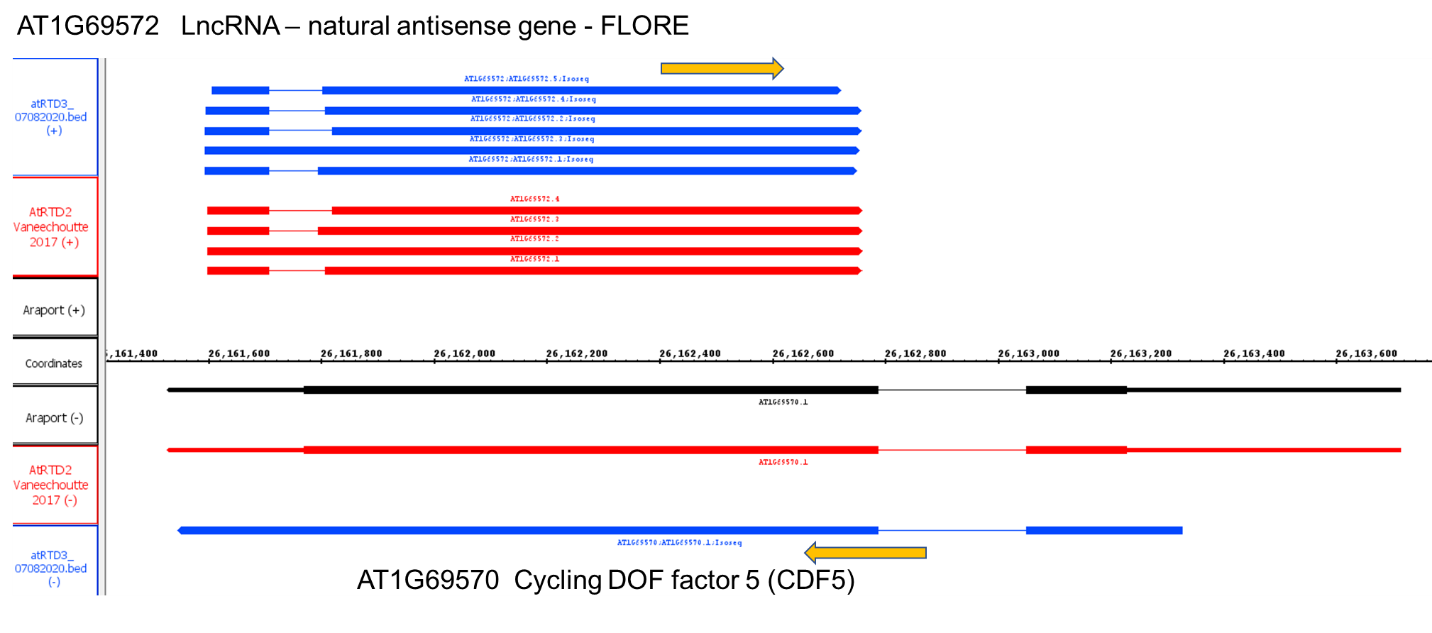

**Fig. S8. Confirmation of AS variant isoforms in the lncRNA, FLORE.** AtRTD3 has five Iso-seq transcripts which confirm differential AS of transcripts from FLORE (Henriques et al., 2017). Transcript structures visualised with IGB are from Araport (black), AtRTD2 (red) and AtRTD3 (blue); arrows show direction of transcription.

A

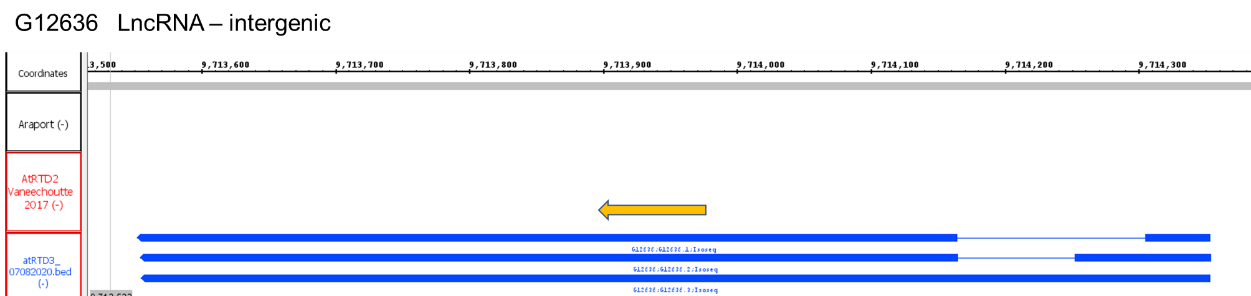

B

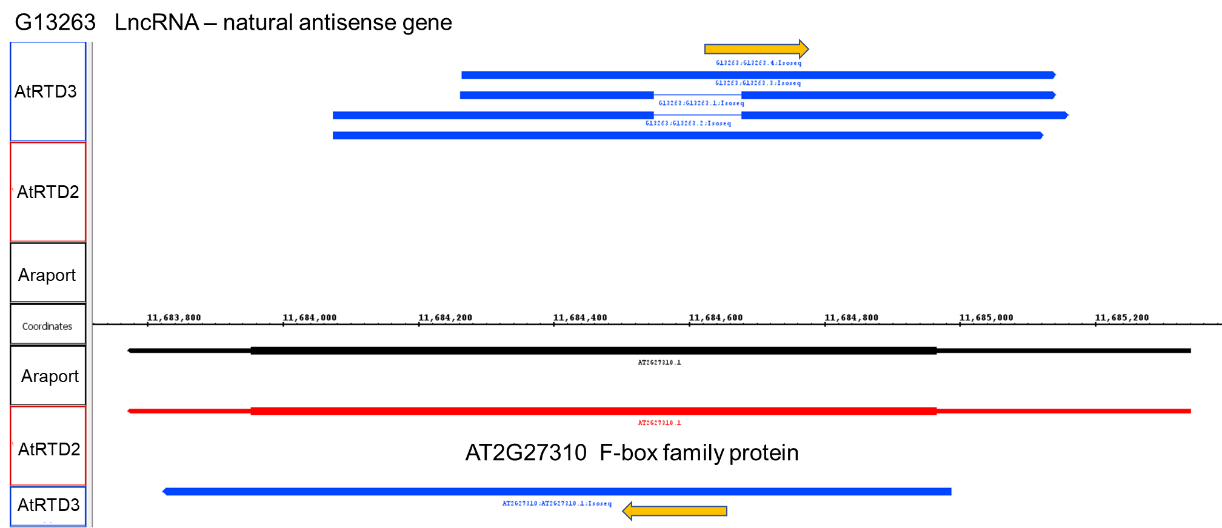

C

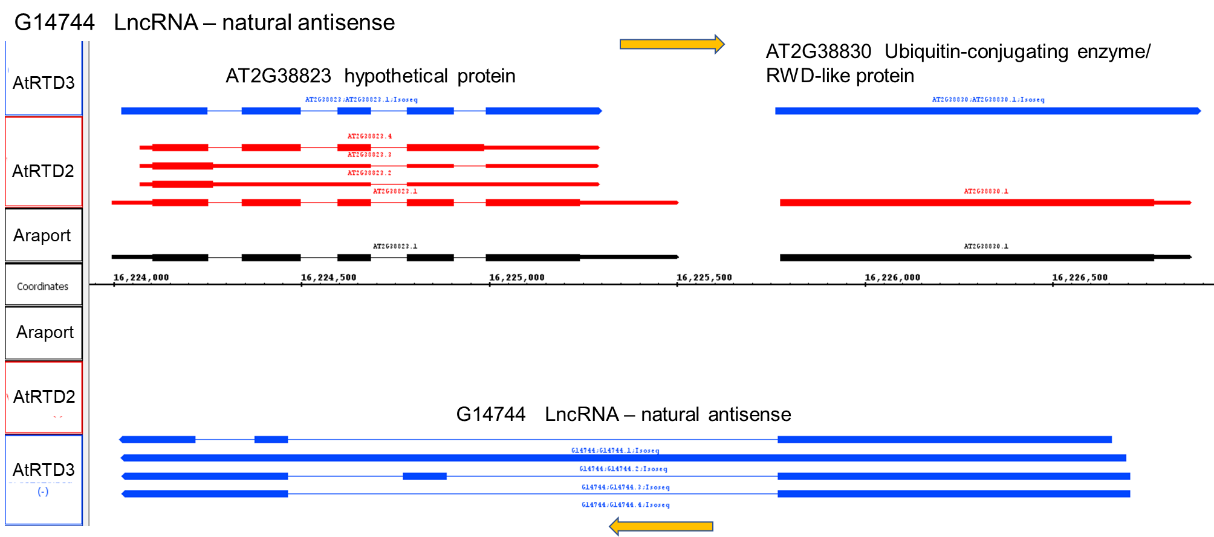

**Fig. S9. Novel genes in AtRTD3 – lncRNAs. A)** G12636 is an intergenic lncRNA gene with three alternatively spliced Iso-seq transcript isoforms; **B)** G13263 has four Iso-seq transcripts with different TSS and spliced and unspliced isoforms; antisense to AT2G27310; **C)** G14744 is antisense lncRNA with four alternatively spliced Iso-seq isoforms which are antisense to two genes, At2G38823 and AT2G38830. Transcript structures visualised with IGB from Araport (black), AtRTD2 (red) and AtRTD3 (blue); arrows show direction of transcription.

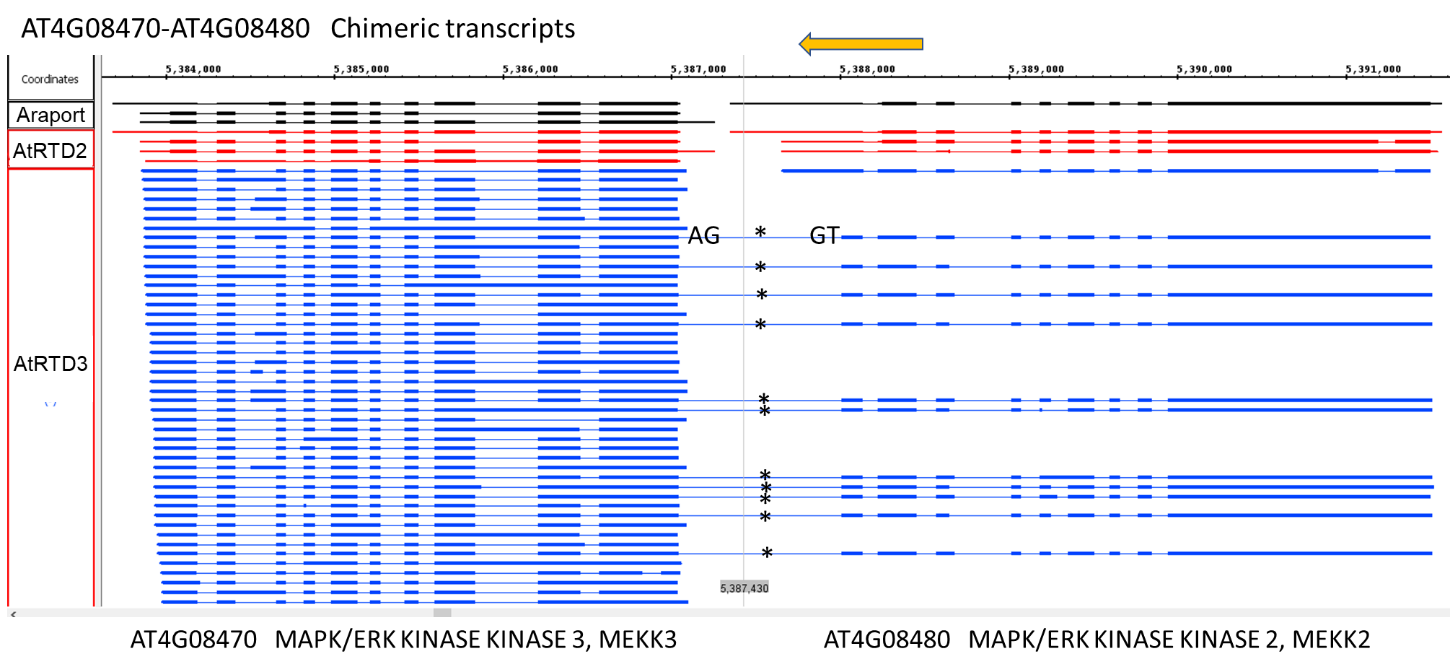

**Fig. S10. Chimeric transcripts.** AtRTD3 contains numerous chimeric transcripts. Two genes encoding MAPK/ERK kinase kinase genes (MEKK3 and 2) generate 11 chimeric transcripts (asterisks) which are linked by an intron. Transcript structures visualised with IGB are from Araport (black), AtRTD2 (red) and AtRTD3 (blue); arrow shows direction of transcription. GT and AG indicate the splice junctions of the intron linking the chimeric transcripts from the 3’UTR of AT4G08480 to the first exon of AT4G08470.

**AT2G29630 – THIAMIN C SYNTHASE (THIC)**

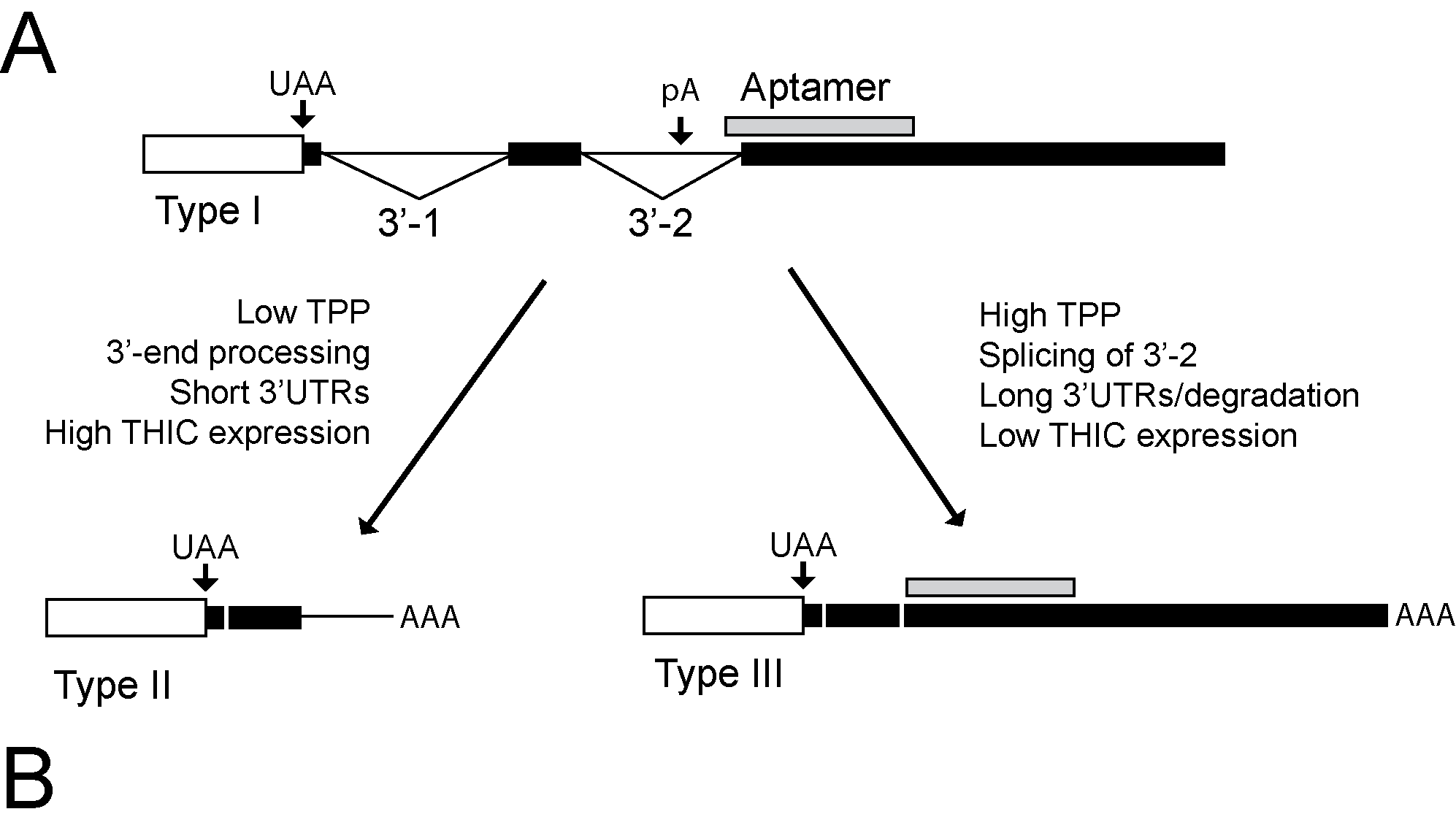

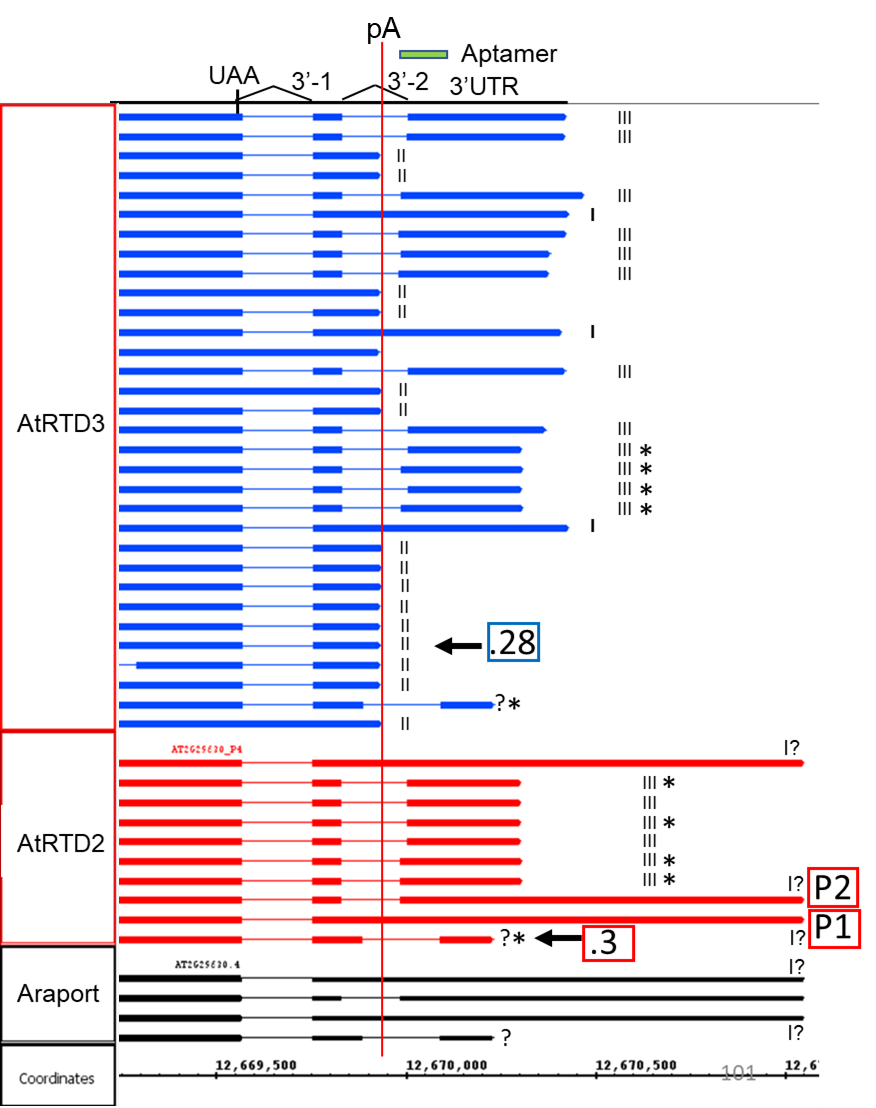

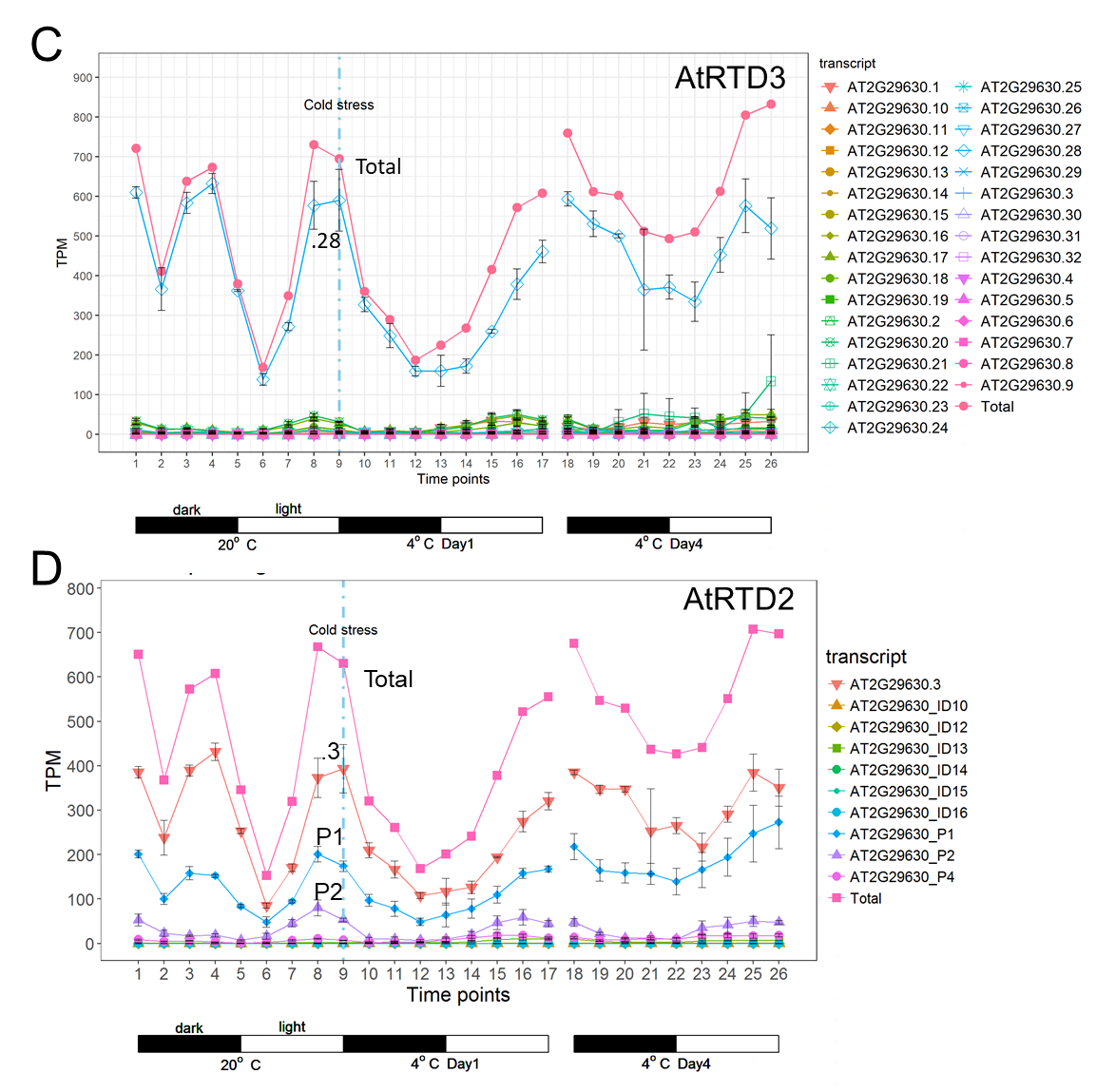

**Fig. S11. Accurate Iso-seq transcript determination identifies THIC RNAs produced by riboswitch. A)** THIAMIN C SYNTHASE THIC (AT2G29630) contains a highly conserved RNA aptamer in the 3’UTR which is bound by TPP to regulate expression via alternative polyadenylation and splicing. Three main RNA classes (Types I, II and III) are produced and type II and III RNAs are generated from the Type I precursor depending on TPP levels. **B)** 3’-ends of transcripts in AtRTD3, AtRTD2 and Araport. Type I, II and III transcripts are clearly observed among the Iso-seq transcripts in AtRTD3. Type II transcripts end at the poly A site in intron 3’-2 (vertical red line). Type III transcripts have longer and variable 3’ends and the 3’-2 intron is removed; type I transcript pre-cursors still contain the 3’-2 intron. Asterisks – incorrectly assembled transcripts in AtRTD2 and Araport. **C)** and **D)** Gene and transcript expression profile in cold treatment time-course with AtRTD3 (**C**) and AtRTD2 (**D**) as reference. Profiles of total expression of the gene are the same but the main transcript in AtRTD3 is a type II RNA (.28) while .3, P1 and P2 transcripts are observed with AtRTD2.

**AT3G17510	CBL-INTERACTING PROTEIN KINASE 1 (CIPK1)**

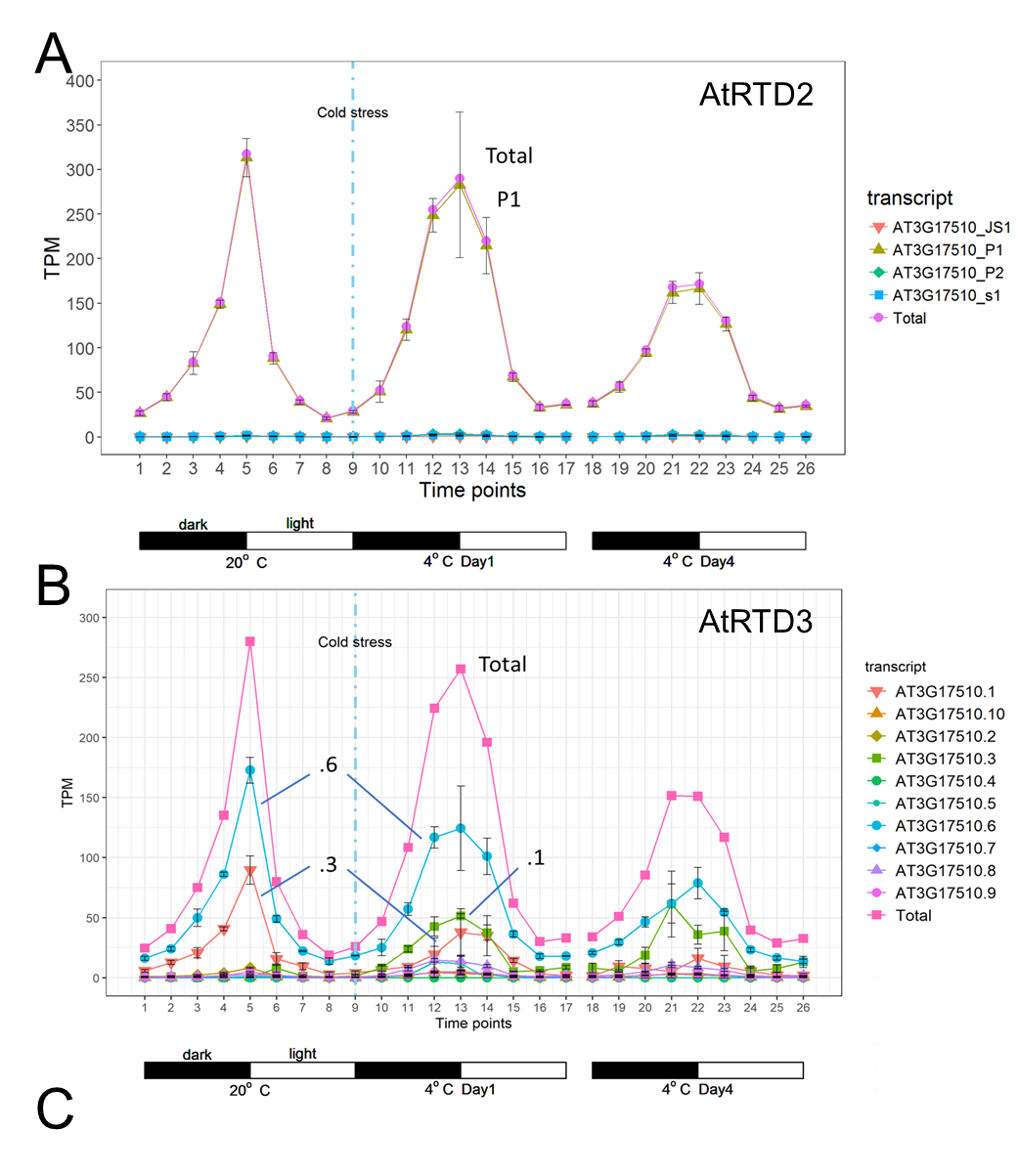

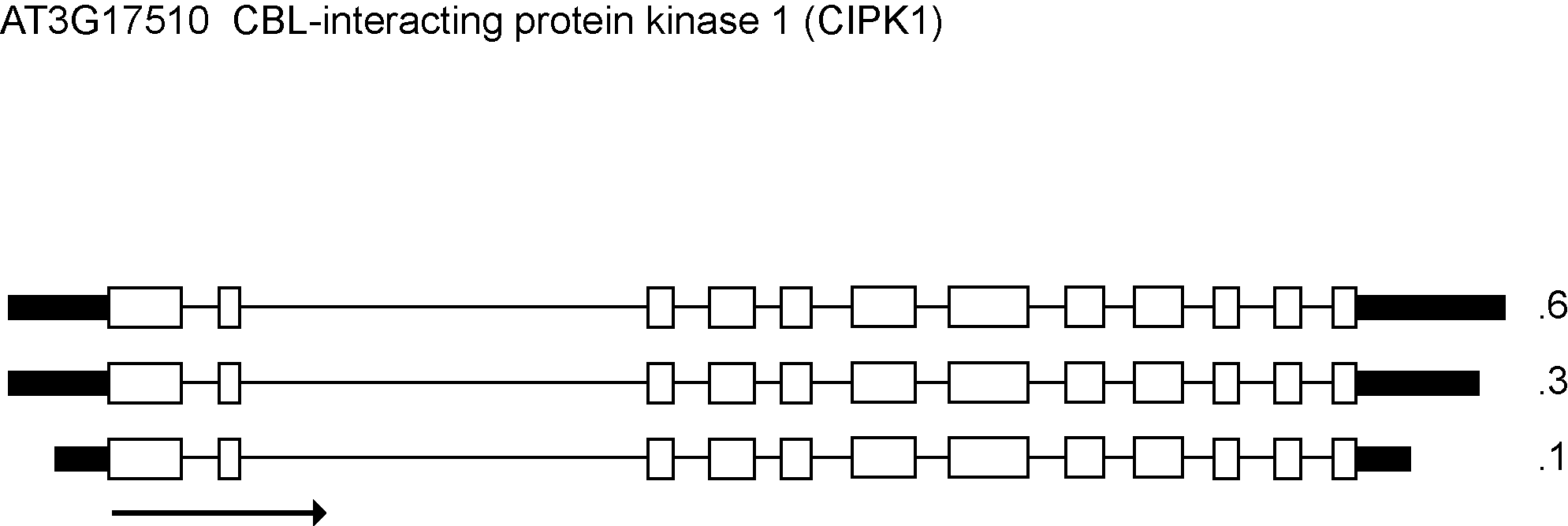

**Fig. S12. Novel cold-induced isoform of CIPK1 from AtRTD3. A)** and **B)** Gene and transcript expression plots of AT3G17510 - CBL-INTERACTING PROTEIN KINASE 1 (CIPK1) using AtRTD2 or AtRTD3 as reference. CIPK has 4 transcripts in AtRTD2 of which only the P1 isoform is highly expressed. Expression is rhythmic, peaking at dawn; low temperature broadens the peak of expression and it reduces with cold exposure. B) AtRTD3 has 10 isoforms of which three (.6, .3 and .1) are the most highly expressed. The .1 isoform is induced by cold. C) All three isoforms code for the same protein; .3 and .6 have the same TSS but different TES; the cold-induced .1 isoform has shorter TSS and TES.

**AT4G25080 - MAGNESIUM-PROTOPORPHYRIN IX METHYLTRANSFERASE (CHLM)**

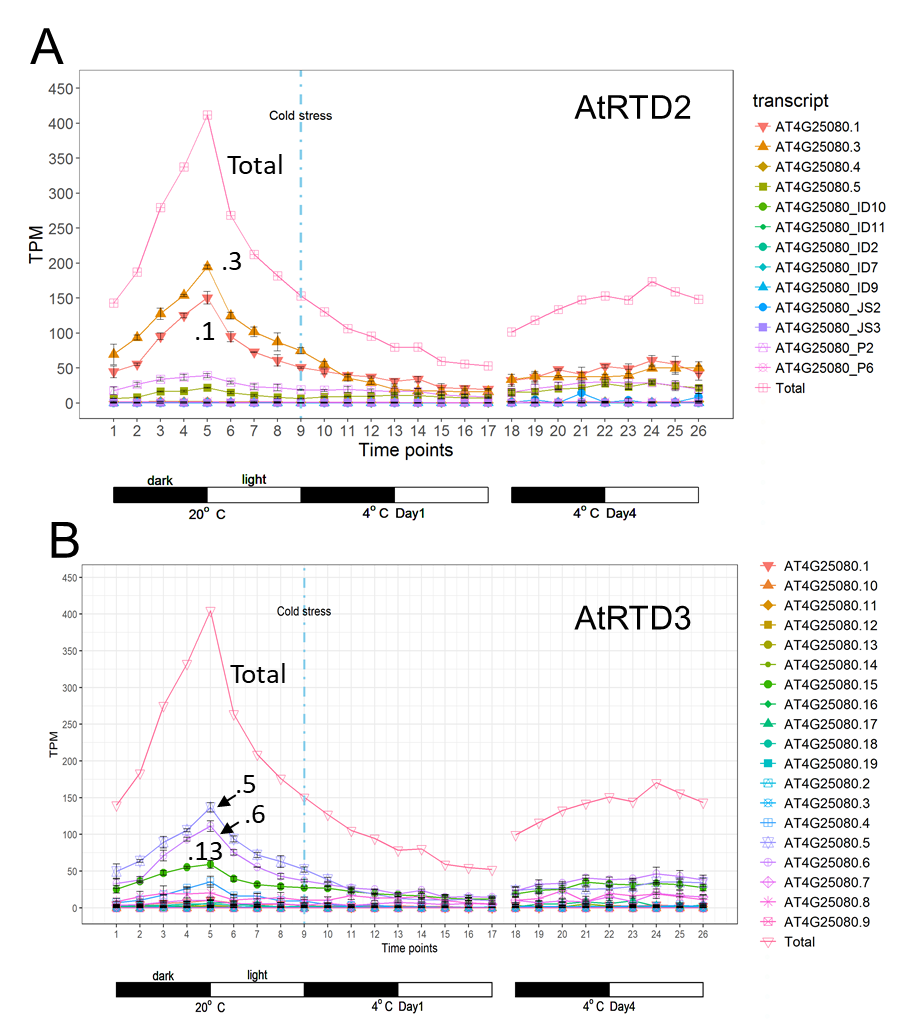

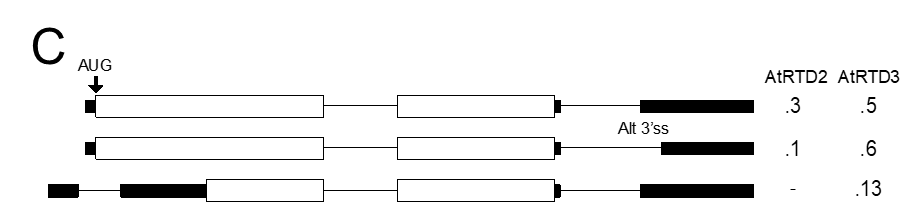

**Fig. S13. Novel transcript isoform in AtRTD3 affects expression levels of main transcripts compared to AtRTD2.** Gene/transcript expression profiles of AT4G25080 using A) AtRTD2 or B) AtRTD3 (lower) reference. A) Two most expressed transcripts in AtRTD2 (.3 and .6) code for the same protein but differ by an alternative 3’ splice site in the 3’UTR (C). B) AtRTD3 identifies a novel expressed transcript (.13) which causes reduction of expression levels of .5 and .6 (equivalent to .3 and .1 in AtRTD2. Total gene expression profiles are the same with both references.
