## Additional file 3 for "A high resolution single molecule sequencing-based Arabidopsis transcriptome using novel methods of Iso-seq analysis"

**Additional File 3 : Comparison of AtIso to AtRTD2 and Araport11**

**for**

**Comparison of AtIso to AtRTD2 and Araport11**

AtIso had lower gene coverage than the short read-derived Araport11 and AtRTD2 transcriptomes [34,45] AtRTD2 used stringent quality filters to remove false spice junctions, redundant transcripts and transcript fragments and contained 82,190 transcripts from 34,212 genes [34]. AtRTD2 incorporated transcripts from protein-coding genes from an early version of Araport11. It also contained significantly increased transcript and alternative splicing diversity compared to TAIR10 and Araport 11 but did not contain any novel gene models compared to Araport11 [34]. Therefore, we compared the genes in AtIso to the current version of Araport11 which contains 38,194 genes with 59,775 transcripts. Of the 38,194 Araport11 genes, 20,663 genes had significant overlapping regions with AtIso genes on the same strand (coverage >50% of total gene length). An additional 719 genes overlapped AtIso transcripts with coverage ≤ 50% of total gene length. Thus, 21,382 (56.0%) Araport11 genes overlapped AtIso genes and 16,812 Araport11 genes (44.0%) had no coverage in AtIso. Of the 21,853 genes in AtIso, 20,194 genes had significant overlapping regions with Araport11 genes on the same strand (coverage >50% of total gene length) with an additional 210 genes overlapping with coverage of ≤ 50% of total gene length). Thus, AtIso contained ca. 1,450 novel genes compared to Araport11. Despite extensive sequencing of a wide variety of tissues and conditions, gene coverage in AtIso was limited to 576% of genomic loci in Araport11.

We next compared transcript identity among the three annotations using TAMA merge to identify transcripts with exactly the same SJs and only differing by <50nt at the 5’ and 3’ ends. There are a total of 209,508 non-redundant transcripts in the three annotations. Only 5,369 (2.56%) transcripts were shared by all three, and 6,167 (2.94%) and 980 (0.4%) transcripts were shared between AtIso and AtRTD2 and AtIso and Araport11, respectively (Fig. S14A) suggesting a high degree of difference among transcripts. Comparison of splice junctions among the three transcriptomes (a total of 183,035 non-redundant SJs) showed that 100,275 (54.78%) SJs were common to all three (Fig. S14B). A quarter of SJs only occurred in AtRTD2 (10,155 - 5.5%); 2,410 (1.3%) were unique to Araport or common to both (33,115 - 18.1%) and 28,035 SJs were unique to AtIso (15%) (Fig. S14B). Thus, there is good agreement of the SJs identified by short and long reads which is in sharp contrast to the small overlaps in transcript identity (Fig. 4B) between long and short read assembled transcripts. The difference is illustrated by only 5.9% of transcripts and 75% of SJs shared between long and short read assemblies. This mainly reflects differences at the start and end positions between long and short read assemblies. Transcript start/end determination is generally inaccurate with short reads and Araport is known to have extensive mis-annotations at 3’ and 5’ UTR regions (mostly over-extended) which were carried over into AtRTD2 [34]. The complementarity between SJs from long and short reads reflects the novel methods of removing false SJs here and in the AtRTD2 assembly [34]. The number of SJs unique to AtRTD2 most likely reflects the higher gene coverage while those unique to AtIso appear to come from long reads discovering SJs of minor isoforms in highly expressed genes. Thus, AtIso contains accurate and diverse transcripts but suffers from poor coverage of around one third of gene regions.

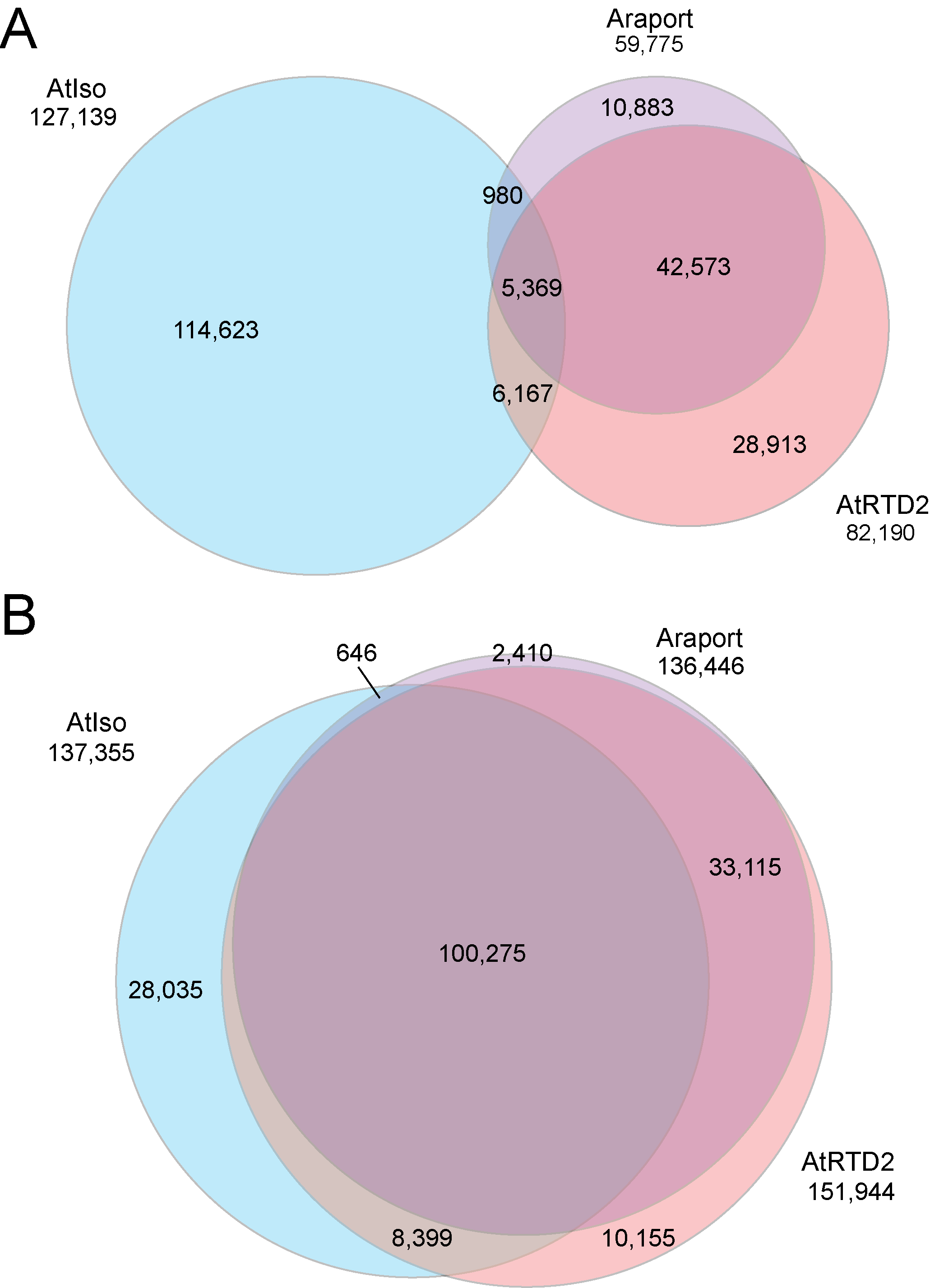

**Figure S14. Comparison of AtIso transcripts and SJs to Araport and AtRTD2. A)** Transcripts; **B)** SJs for AtISO (light blue), Araport (lilac) and AtRTD2 (pink).
